# Vision dominates sound in mediating classic cue-induced microsaccadic eye movement modulations in rhesus macaque monkeys

**DOI:** 10.1101/2025.09.25.678668

**Authors:** Tatiana Malevich, Matthias P. Baumann, Yue Yu, Ziad M. Hafed

## Abstract

Visual onsets in the environment cause a short-latency reduction in saccade generation likelihood. Concomitant with that, the metrics of the saccades that do happen near the time of saccadic inhibition reflect the landscape of stimulus-driven neural responses in topographically organized brain maps. While certain aspects of saccadic inhibition have already been explored from a neurophysiological perspective, several questions remain, including which sensory-motor pathways propagate visual input information to the final oculomotor control circuitry with such short latencies. Here, motivated by evidence of a dissociation between temporal and spatial aspects of saccadic inhibition, we devised behavioral paradigms experimentally manipulating either aspect. Rhesus macaque monkeys maintained gaze fixation, and we characterized visually-driven microsaccadic inhibition with and without paired auditory pulses. In one condition, the sound pulses were spatially-nonlateralized, thus only affecting the temporal processes associated with microsaccadic inhibition; in another, the pulses were spatially-lateralized, affecting both the temporal and spatial components. We found that sound pulses alone barely affected microsaccade rates. However, when paired with visual stimuli, they accelerated the timing of visually-driven microsaccadic inhibition by ∼14-18 ms. In terms of microsaccade directions, spatially-nonlateralized sounds interfered with, but did not eliminate, visually-driven direction modulations. Surprisingly, lateralized sounds had a much weaker effect on microsaccade direction modulations, even when the sound source was in the opposite hemifield as the appearing visual stimulus. We conclude that vision dominates sound in mediating classic microsaccadic inhibition, and we argue that our paradigms are ideal for neurophysiological investigations of the parallel sensory-motor pathways contributing to such inhibition.

**Significance:** Coordination of internal oculomotor eye movement plans with exogenous sensory events entails rapid inhibition of the saccadic eye movement system, but the pathways mediating the inhibition are unknown. We devised novel behavioral paradigms in primates, involving multisensory stimulation, which allow neurophysiological experiments to isolate the contributions of individual sensory, motor, and sensory-motor brain areas to saccadic inhibition. Such paradigms underscore that inhibition involves distinct temporal and spatial processes that are dominated by the visual modality.

## Introduction

Sensory transients in the environment have a powerful, unavoidable impact on the behavior of the oculomotor system, giving us the subjective impression of reflexively orienting towards such transients with very short latencies. However, there is an even earlier behavioral outcome of sensory transients, and one that precedes reflexive looking (1). This earlier outcome manifests itself as a robust, short-latency modulation not only of saccades (2–5) and microsaccades (6–9), but also of slow ocular position drifts during fixation, which are on the scale of approximately 1 min arc eye movements (10, 11), as well as smooth pursuit (12–14).

In saccades and microsaccades, our focus here, sensory transients cause an almost complete cessation of eye movement generation, in a phenomenon called saccadic (or microsaccadic) inhibition (1–4, 6). While parts of the underlying neural mechanisms for saccadic inhibition have been explored (15, 16), much remains to be learned (1, 17), especially given the plethora of ways in which sensory information from the periphery can find its way to the final oculomotor control pathways. Prior computational accounts have supported the idea of a potential anatomical dissociation between temporal and spatial factors mediating saccadic inhibition (18, 19), meaning that the same sensory transient can simultaneously activate both a temporal inhibition signal for altering saccade frequencies as well as a spatial biasing signal for influencing eye movement metrics and kinematics. While this is consistent with behavioral and neurophysiological evidence (16–18, 20–23), experimental tests of the relative contributions of cortical and subcortical brain circuits to saccadic inhibition require paradigms that can manipulate the inhibition’s temporal and spatial factors. Here, we explored and characterized such paradigms.

In the visual modality, manipulations of the spatial extent of a visual stimulus, as well as the spatial configuration of multiple simultaneously appearing ones, have been particularly informative about the spatial component of saccadic inhibition (18). These results were also significant because they added to growing evidence (8, 9, 15, 16, 24–28) that saccadic inhibition can be effectively studied in an ideal animal model for neurophysiological mechanisms, the rhesus macaque monkey. Having said that, there is a need for additional ways to explore the temporal and spatial components of saccadic inhibition. For example, a visual stimulus alone not only manipulates the spatial aspect (via biasing of spatial maps representing the sensory environment), but the transient visual onset itself simultaneously jumpstarts the temporal processes. Here, we used multisensory stimulation to allow a further experimental differentiation. We paired a visual stimulus onset in either hemifield with either a spatially-nonlateralized sound pulse relative to the vertical meridian (thus only affecting the temporal processes of the inhibition) or a spatially-lateralized one that was biased to one visual hemifield more than the other (thus affecting both the spatial and temporal processes). Importantly, we performed our experiments in monkeys, as an obligatory prerequisite for allowing subsequent neurophysiological experiments in the sensory-motor pathways of the same animals.

In what follows, we describe the relative contributions of sound and vision to microsaccadic inhibition in rhesus macaque monkeys. We show that the visual modality dominates effects on both microsaccade rates and directions, and that simultaneous auditory stimulation accelerates and interacts with the visually-driven modulations. We believe that our experimental paradigms and analyses are a useful tool for neurophysiological investigations of multiple sensory-motor pathways mediating a highly robust oculomotor phenomenon.

## Methods

### Experimental animals and ethical approvals

We collected eye movement data from two adult, male rhesus macaque monkeys (F and A, aged 14 and 12 years, and weighing 13 and 9.5 kg, respectively). The monkeys were prepared for behavioral and neurophysiological experiments previously (e.g. (29, 30)). Briefly, each monkey was implanted with a head-holder for stabilizing head position during data collection, as well as with a scleral search coil in one eye, allowing accurate and precise eye tracking using the magnetic induction technique (31, 32).

All experiments were approved by ethical committees at the regional governmental offices of the city of Tübingen (Regierungspräsidium Tübingen).

### Laboratory setup

The monkeys sat in a dark room in front of a CRT display having a gray background of 26.11 cd/m^2^ luminance (Fig. 1A). The display was centered at eye height, and it was 72 cm in front of the animals. The display subtended ∼30 deg horizontally and ∼23 deg vertically.

**Figure 1.**
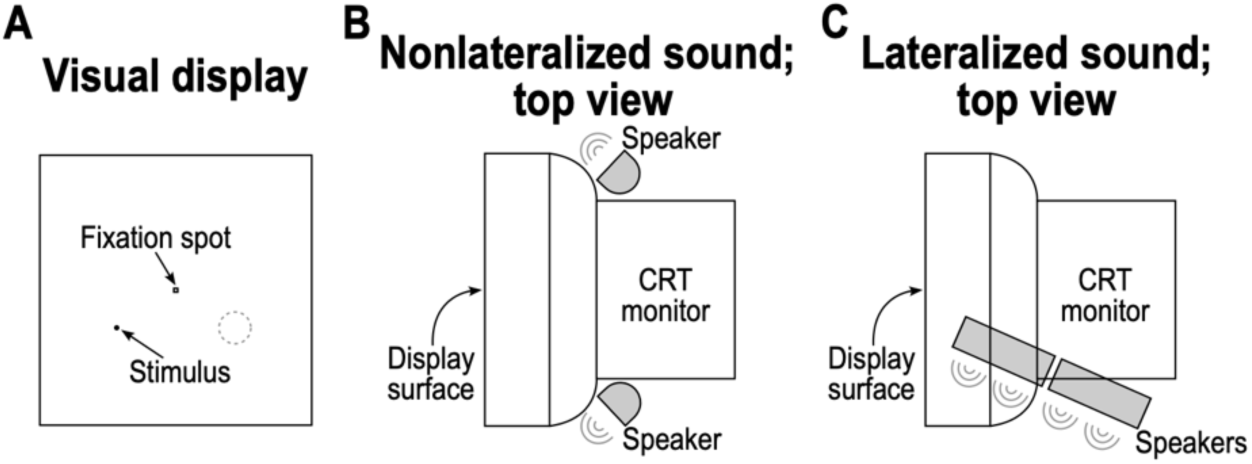
Physical configuration of the visual and sound sources in the experiments presented in this study. **(A)** The monkeys fixated a central fixation spot, and we presented a visual stimulus onset in the lower visual field. The stimulus could appear either to the left or right of the fixation spot in terms of its horizontal position; the shown trial is one with a spot appearing to the left of the fixation spot, and the dashed circle (which was not shown on the display) indicates that, in other trials, the stimulus could appear in a mirror-symmetric location. **(B)** In the spatially-nonlateralized condition, we placed two speakers behind the visual display, as shown relative to the CRT monitor that was presenting visual stimuli (Methods). The speakers were on the same table holding the monitor, and the sound source was thus also in the lower visual field relative to the monkey. **(C)** In the spatially-lateralized condition, we placed two speakers below the CRT monitor, and in a position to emit sound in one direction (the shown example is for the sound being emitted towards the right of the monkey; in other blocks of trials, the speakers were moved to a mirror-symmetric position, such that they emitted sounds to the left side of the monkey; Methods).

For trials with a sound pulse (see experimental procedures below for more details), we used a pair of audio speakers that were placed in two possible configurations (Fig. 1B, C). For the spatially-nonlateralized configuration, the goal was to have a diffuse sound source that would not be spatially biased to either the right or left visual hemifield of the monkey (when it was looking directly at the display center). Thus, each speaker was placed on the same table holding the display, but with one speaker being behind the bottom right corner of the display (behind by 22 cm from the front of the display and 21 cm below the center of the display) and the other speaker being behind the bottom left corner of the display (again, behind by 22 cm from the front and 21 cm below the center; Fig. 1B). Each speaker was 9 cm wide and 15 cm high, and it was placed upright. The right speaker was aimed towards and to the right of the monkey (that is, diagonally), and the left speaker was again aimed diagonally, but now towards and to the left of the monkey. This way, the sound surrounded the display in a balanced manner and was not spatially lateralized to either the right or left visual hemifield. For practical considerations in our laboratory environment, we had no space to place the two speakers directly behind the display, to give rise to a centered sound source.

For the spatially-lateralized sound configuration, we again placed two speakers (each 9.5 cm wide and 24.5 cm high), but this time, they were placed directly next to each other in either the right or left visual hemifield of the monkey (Fig. 1C). Specifically, the two speakers were laid down on their side on a platform directly under the table holding the display. Thus, the two speakers formed a long, continuous, and horizontal line of sound source (49 cm long in total). This horizontally-lying sound source was aimed diagonally towards and to one side of the monkey, in order to bias the sound waves towards one visual hemifield. For example, if the speakers were placed under the right half of the display (under by 22 cm from the display’s bottom), then they were oriented such that they emitted sound diagonally towards and to the right of the monkey. If they were placed under the left half of the display, they were oriented such that they emitted sound diagonally towards and to the left side of the monkey. Thus, the emitted sound pulse was biased to one visual hemifield when the monkey was fixating the display center, but it could, of course, reach both ears since the speakers were in the environment and not directly placed inside each ear. This was fine for our purposes because it was sufficient to introduce a spatial lateralization in the sound source (33). Note that this placement of the speakers in this case also made the sound source in these experiments similar to the experiments with the spatially-nonlateralized sounds (i.e. still in the lower visual field like the visual stimuli).

We measured the sound intensity at the position of the monkeys’ heads using a sound meter (NL-52A, RION, Tokyo, Japan). For monkey F, the sound intensity at the head was 69-89 decibels (db) in the spatially-nonlateralized configuration (89 db was only used in the first session) and 76 db in the spatially-lateralized configuration. For monkey A, the sound intensity at the head was 89 db in the spatially-nonlateralized configuration, and it was 76 db in the spatially-lateralized configuration. The slight difference in intensities between the lateralized and nonlateralized conditions was a function of the different physical setup and the different speakers used in the two sets of experiments. The background noise level was 51-54 db; this constituted just the ambient noise level of the laboratory environment, and it was continuously present. In all cases, the sound pulse that we used was a pure tone (1 kilohertz, kHz, in both monkeys, except for two sessions in monkey A, in which we used 2 kHz) of 50 ms duration. Thus, our sound pulses were clearly audible (suprathreshold), which was our intent, but they were not so loud as to cause startle reflexes by the monkeys (like blinking); in fact, monkey F was initially startled by the intensity used for monkey A, and that is why we reduced the sound intensity (from 89 db) for this monkey after the first session (however, for the later sessions in which we tested 69 db and 79 db intensities, we did not see any differences in the properties of saccadic inhibition). In any case, at least in humans, microsaccade rates were shown to not be affected by the emotional arousal level of sound stimuli (34). As for the choice of sound frequency, it was subjectively chosen by us, to again allow audibility but without discomfort that might be associated with very high pitch.

Other aspects of the experimental setup (such as visual display control) were the same as those described previously (29, 30, 35, 36). Briefly, we controlled the experiments using a custom-made computer system based on PLDAPS (37) and the Psychophysics Toolbox (PTB-3) (38–40). The Psychophysics Toolbox allows querying the graphics system of the display computer at every frame refresh, thus allowing frame-by-frame display control. We also used Psychophysics Toolbox functions for triggering sound pulses with a temporal precision of a few milliseconds. Specifically, the same toolbox has a high precision sound driver, which is initialized with the command “InitializePsychSound” at the beginning of the experiment. Then, at the same time as when we requested the refresh of the frame for the visual stimulus onset in a given trial, we called the command “PsychPortAudio”. This command drove the speakers to emit the tone. We estimated a deviation of about +/-5 ms between the sounds and visual onsets, given the refresh rate of our display. This temporal precision (on trials in which we paired the sound pulse with a visual stimulus onset) was sufficient for remaining within the expected temporal window in which auditory-visual multisensory integration could happen in the brain (41–43). We recorded all data (at 1 kHz sampling resolution) using the OmniPlex data acquisition system from Plexon, and our display’s refresh rate was 85 Hz.

### Experimental procedures

The behavioral task for the monkeys always consisted of maintained gaze fixation on a central, black fixation spot of 10.9 by 10.9 min arc dimensions (on an otherwise gray background; Fig. 1A); our approach was to analyze the properties of microsaccades occurring during such gaze fixation.

#### Spatially-nonlateralized sound task

In one set of blocks and sessions, we pseudorandomly interleaved trials of three types: control trials with only the onset of a spatially-nonlateralized sound pulse (20% of all trials within a session), visual-only trials in which only a visual onset occurred without a sound pulse (40% of all trials), and visual plus auditory trials in which the visual onset was paired with a simultaneous spatially-nonlateralized sound pulse (40% of all trials within a session). In the visual onset trials (whether with or without the spatially-nonlateralized sound pulse), the stimulus consisted of a small black spot (100% Weber contrast) of 0.36 deg diameter, and it remained on the display for 100 ms. Trials ended 500-800 ms after the offset of the visual stimulus (or 500-800 ms after the onset of the sound pulse on the control trials), and the monkeys were rewarded for maintaining fixation on the central fixation spot until trial end. The visual stimulus was placed at an eccentricity of 0.5-12 deg from the central spot, and in the lower visual field (1-8.5 deg direction from the horizontal). This placement, which was constant within a block of trials, was dictated by where we might encounter response fields in areas like the primary visual cortex and superior colliculus in subsequent neurophysiological investigations with our paradigm. Moreover, lower visual field stimuli were closer to the sound sources, allowing us to amplify the potential impacts of either spatially lateralized or nonlateralized sounds on visually-driven microsaccadic modulations. We collected 171-1489 trials per session in each monkey, and we included data in this study from 10 sessions in monkey F (7180 trials in total) and 40 sessions in monkey A (22403 trials in total).

Note that our visual stimulus duration was long enough to ensure observing saccadic inhibition in monkeys, with such inhibition typically starting within less than 100 ms from stimulus onset (8, 18, 23, 24). In fact, much briefer flashes can still elicit robust saccadic inhibition in these animals (9). Moreover, we made sure that our sound pulse was sufficiently long (50 ms) to have sufficient overlap in the onset times between the visual and auditory stimuli, even if there were a few milliseconds of timing jitter from the hardware.

In the data analyses for this task, we also included visual-only trials from an additional set of blocks and trials from a different behavioral paradigm. In this paradigm, we were varying the visual properties of the stimuli across trials, such as the Weber contrast and luminance polarity, for purposes not directly related to the current study. However, the 100% contrast, black condition in that other paradigm was identical to our visual-only trials here. Thus, we took these particular trials (32%) and collapsed them with the visual-only trials described above. By doing so, we obtained an additional 2370 trials over 10 sessions in monkey F and 4378 trials over 20 sessions in monkey A.

#### Spatially-lateralized sound task

We ran separate blocks of trials in which the sound pulses were spatially-lateralized. In these blocks and in monkey A, we interleaved two trial types within a given session: control trials with only the onset of a spatially-lateralized sound pulse (∼33.3% of all trials within a session), and vision plus sound trials in which the visual onset was paired with a spatially-lateralized sound pulse (∼66.6% of all trials). In monkey F, these types of trials were also interleaved with visual-only trials, resulting in the following trial ratio within a given session: control trials with only the onset of a spatially-lateralized sound pulse (20% of all trials), vision plus sound trials (40% of all trials), and visual-only trials (40% of all trials). For the later analyses, we collapsed the visual-only trials from this task in monkey F (5648 trials) with the visual-only trials from the spatially-nonlateralized sound task described above.

We typically ran one block with the sound source on the right, and one block with the sound source on the left on each day (the order of the right- and left-sound blocks was counterbalanced across sessions), and we collected 223-1022 trials per sound location session per monkey. In the analyses, we included data from 11 sessions in monkey F and 18 sessions in monkey A, giving a total of 14126 and 18001 analyzed trials for each monkey, respectively.

In most days, we ran the spatially-nonlateralized sound blocks described above first, and then two spatially-lateralized sound blocks (one for a right and one for a left sound source). However, for some sessions, we sometimes only ran the spatially-nonlateralized sound blocks during each session, and sometimes only the spatially-lateralized sound blocks. This approach was not a problem for our analyses because microsaccadic inhibition is a stationary process across many trials, blocks, and sessions. For example, it is not altered across thousands of trials in individual monkeys (8) (also see Figs. 2, 4, 7 in Results), as well as in humans (44). Also, in neurophysiological experiments, one is very likely to block the presentation of stimuli like this, and experimental constraints can sometimes force a reordering of which tasks to run first, and so on.

**Figure 2.**
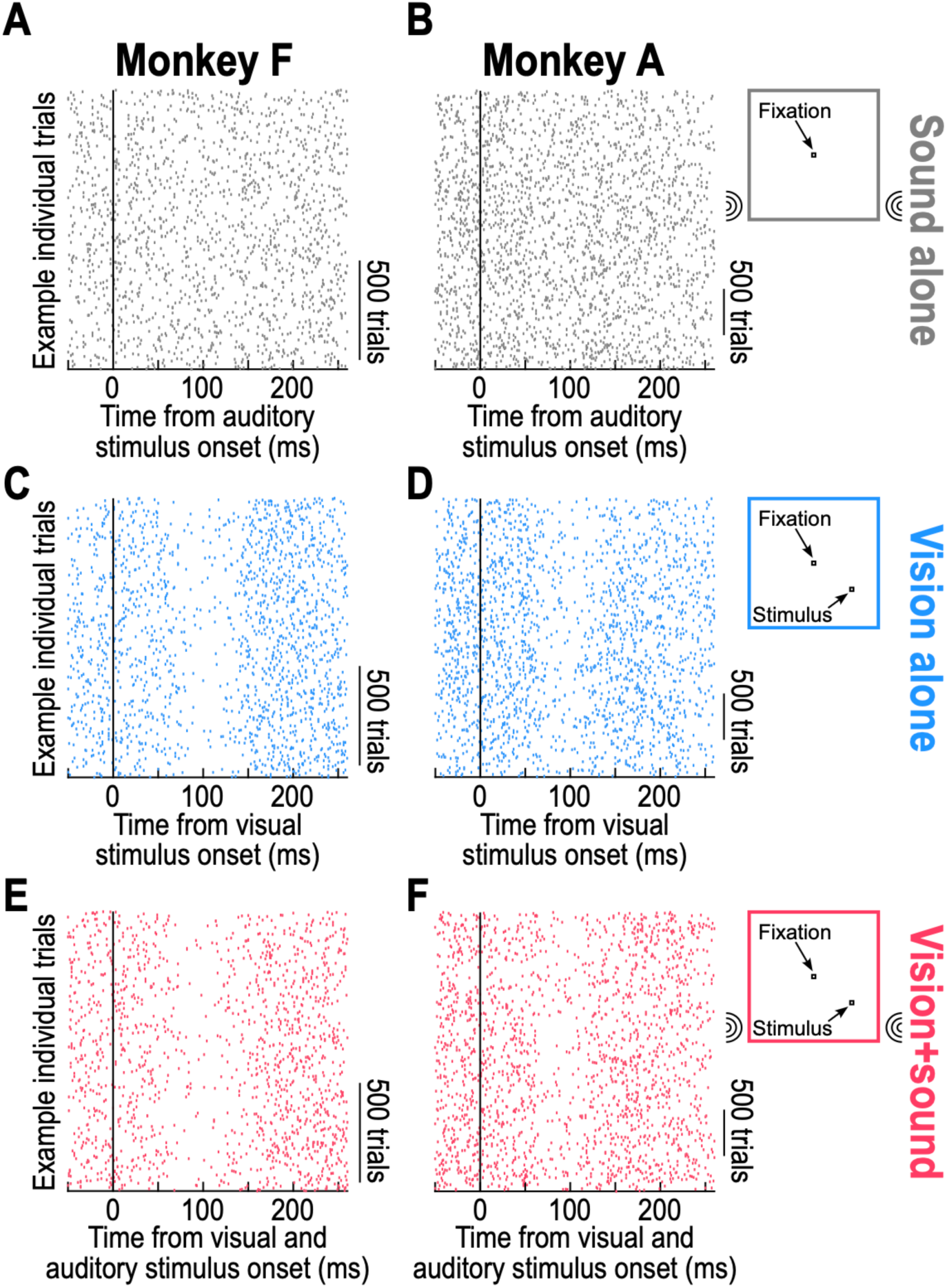
Spatially-nonlateralized sound alone does not cause clear microsaccadic inhibition in rhesus macaque monkeys. **(A, B)** For each of monkey F (**A**) and monkey A (**B**), we plotted a raster of microsaccade onset times relative to the onset of a brief, spatially-nonlateralized sound pulse (Methods). The visual display did not contain any transient stimuli during these trials, and the monkeys were instructed to simply fixate the stable, central fixation spot. Each row is an individual trial, and each dot indicates a microsaccade onset time. There were barely any changes in microsaccade likelihood after the occurrence of the sound tone (also see Fig. 3 for more detailed quantification). **(C, D)** In sharp contrast, a visual stimulus onset (Methods) caused robust microsaccadic inhibition (<100 ms after visual stimulus onset) in both monkeys, as expected in these animals (1, 8, 9, 17, 18, 20, 24–26, 28). **(E, F)** Such robust microsaccadic inhibition also occurred when the visual stimulus onset was paired with a simultaneous sound pulse identical to that used in **A**, **B**. Note that the inhibition in this case started slightly earlier than in **C**, **D**, as quantified more clearly in Fig. 3. Also note that we collected more trials in each monkey for all subsequent analyses (Methods), meaning that the shown trials here are from example subsets of individual trials (the shown numbers of trials can be inferred from the vertical scale bars).

Note that our aim in this study was not to perfectly match the location of the visual stimulus with the location of the spatially-lateralized sound source. Rather, we wanted to investigate how hemifield biases in the sounds influenced visually-driven microsaccadic modulations when the visual stimulus was in a congruent or incongruent hemifield. Since our visual stimulus was just a single spot at one location, our goal was achieved with our audio speaker placements. Specifically, because it is expected that the microsaccades would be directed towards the visual field stimulus location shortly after its onset (18), having the sound in the same or opposite hemifield was sufficient to explore the impacts of the sounds on these modulations (there was no other competing visual stimulus in the hemifield, other than the experimentally-presented spot). Moreover, auditory receptive fields in relevant multisensory structures for our task, such as the superior colliculus, are quite large, spanning from 45 to 100 deg in diameter in the periphery (42), and multisensory integration was shown to be tolerant to small spatial disparities between unisensory stimuli (45). Finally, there are successful examples in the literature, in which the effects of multisensory integration on neuronal activity (46) and on saccadic behavior (47) were achieved with stimuli matched by hemifields.

### Data analysis

Data analysis was performed in Matlab 2020b (MathWorks, USA) using custom scripts.

We detected all microsaccades using our previously established methods (48, 49), and we removed all trials with blinks (especially the case in monkey F’s first session with 89 db sound pulses). We included all trials in which the monkeys were rewarded in our analyses, and we characterized the properties of microsaccadic inhibition. For any condition, we pooled data from all sessions in which the condition was run, again because microsaccadic inhibition in monkeys is a highly stationary process across thousands of trials (8).

To compute microsaccade rate, we used a moving window approach. For every trial, we stepped a window of 50 ms width in steps of 2 ms, covering a range of times around stimulus (visual, auditory, or both) onset. We then counted the number of microsaccade onsets that occurred within this time window and divided by 50 ms to get a rate estimate (movements per second). We then averaged all rate curves from all trials of a given type to get an average microsaccade rate curve, and we estimated the standard error of the mean for each time sample across trials.

For some visualizations of microsaccade rate curves, we applied a baseline correction, in order to make the different curves being visualized have the same starting level at stimulus onset (independent of the actual underlying rate in terms of movements per second). To do so, we computed the average microsaccade rate curve of a given condition. Then, we took the average rate value of this curve in the final 50 ms before stimulus onset, and we divided the entire rate curve by this baseline measurement. Thus, the normalized rate curve started at a value near 1 regardless of the true underlying prestimulus microsaccade rate.

To measure microsaccade direction modulations, we first classified a movement’s direction according to the hemifield towards which it was directed. Because microsaccades in monkeys orient themselves towards the direction of the appearing stimulus with a relatively high angular fidelity (8, 18, 50), and because we had only one single visual stimulus location in any given trial, such hemifield classification was sufficient to reveal microsaccade direction modulations associated with the visual onset events. A similar situation was true when there was only a spatially-lateralized sound pulse in a given trial. In fact, even in the vision plus sound trials, we had the visual stimulus appear in the lower visual field, consistent with the location of the sound pulses (thus, meaning that the hemifield direction classification was all that we needed). After classifying individual microsaccade directions, we used two methods for assessing direction modulations after stimulus onset.

In the first method, we repeated what we had done previously in monkeys (e.g. (9)). Specifically, for each time window (of 50 ms width; stepped in steps of 2 ms), we calculated the fraction of all occurring microsaccades within the time window that were directed towards the stimulus hemifield (or towards the right or left hemifield in some analyses). Thus, we measured microsaccade direction biases independent of rate modulation (18). If the calculated fraction was 0.5, then microsaccades were equally likely to be in one or the other direction; if it was higher, then microsaccades were more likely to be towards than opposite the stimulus hemifield.

In the second method, we created a metric called the baseline-subtracted rate difference. Here, we were motivated by the observation that individual subjects can have intrinsic idiosyncratic biases in microsaccade directions before any stimulus onset. Thus, for one hemifield or the other, the “fraction towards” method might show a value deviating from 0.5 even in prestimulus intervals. This can complicate looking for stimulus-induced changes in such biases. Thus, we performed baseline subtraction. For a given stimulus, we calculated microsaccade rate curves exactly as above, but now separately for only microsaccades towards or opposite the visual hemifield of a stimulus (or rightward and leftward microsaccades, independent of stimulus location, in some analyses). Then, for each such curve, we measured average microsaccade rate in a prestimulus interval (-50 ms to -1 ms from stimulus onset), and we subtracted this prestimulus value from the rate calculations in each time sample of the rate curve. This centered the rate curve for towards microsaccades around zero movements per second, and it centered the rate curve for opposite microsaccades again around zero movements per second. We then took the difference between the baseline-subtracted towards and opposite rate curves to identify stimulus driven direction modulations. In this case, the metric starts at around zero before stimulus onset, and it is positive if stimulus onset caused an increased bias towards the stimulus location (independent of whether the monkey already had an initial bias in this direction or in the opposite direction).

We also quantified the timing of microsaccadic inhibition. To do so, we created a metric that we called L_B25_, which measured the time at which the microsaccade rate curve after stimulus onset dropped by 25% relative to the prestimulus rate. This metric can help characterize whether microsaccadic inhibition is accelerated or delayed by multisensory stimulation. To calculate it, we took the average microsaccade rate curve of a condition. Then, we measured the average prestimulus microsaccade rate in an interval from -50 ms to -1 ms from stimulus onset. Then, we calculated the 75% value of this baseline rate. We marched forward in time from stimulus onset until microsaccade rate first dropped to this 75% point. This time was defined as the latency to the 25% modulation in microsaccade rate (L_B25_). Note that we picked 25% because it gave us a robust timing value; we were less interested in the absolute onset time of microsaccadic inhibition than in actually comparing this timing across multi- and unisensory stimulation conditions. As a result, we picked a level for which we were certain that microsaccadic inhibition had indeed occurred.

In all conditions, we had thousands of repetitions per monkey (see above and Figs. 2, 7 for examples). Thus, we were highly confident that our reported means were very representative of true population means, and we used descriptive statistics (mean and SEM ranges) throughout the figures of the study. We also used inferential statistics to assess the effects of our manipulations on microsaccadic inhibition latencies. Specifically, we performed nonparametric permutation tests on the differences in the obtained L_B25_ metrics. This approach enabled statistical inference despite having only a single L_B25_ value per condition. Namely, for each comparison, we pooled trials from the two conditions under consideration into a single dataset and randomly reassigned their labels while preserving the original ratio of trial counts across conditions. We repeated this procedure 10000 times. On each iteration, we computed the average microsaccade rate for each condition, derived the corresponding L_B25_ value, and calculated the difference in L_B25_ between the two conditions. The experimental L_B25_ difference was then added to the resulting permuted distribution, and statistical significance was assessed by computing Monte Carlo *p*-values. These were defined as the proportion of permuted differences (under the null hypothesis) that were at least as extreme as the experimentally observed effect. A critical α level of 0.05 was used to determine significance. Multiple comparisons were corrected, when necessary, using the step-down Bonferroni-Holm method (51); the corresponding *p*-values are reported as *p_BH_* values. The provided 95% confidence intervals (*CI*s) were computed as the 2.5^th^ and 97.5^th^ percentiles of the permuted (i.e. null) distribution (that is, observations outside these intervals were considered statistically significant).

To assess the effect of multisensory stimulation on the magnitude of microsaccadic inhibition, we repeated the procedure described above, but this time generating permuted distributions of the differences between the minimum microsaccade rates (i.e. maximal inhibition values) in the poststimulus interval (from +1 to +150 ms relative to stimulus onset). All other steps of the permutation tests remained the same.

Finally, to assess potential adaptation effects of microsaccadic inhibition, especially for sound-only conditions, we reanalyzed the microsaccade rates but now as a function of trial history. In a first analysis, we picked sound-only trials for which the previous trial was either a visual-only trial or a trial with a sound pulse (sound-only or vision plus sound trials). We then plotted microsaccade rates for these two groups of trials separately. If prior exposure to a sound causes adaptation in the present trial, and thus reduces microsaccadic inhibition, then the stimulus-driven rate modulations in the two groups of trials should have looked different from each other. If, on the other hand, microsaccadic inhibition did not adapt, then the sensory exposure of the previous trial should not have influenced microsaccadic inhibition. In a second analysis, we looked for more long-term effects of sensory exposure. For each condition (visual-only, sound-only, or vision plus sound), we grouped trials according to whether they occurred in either the first third of all trials within a given session or the final third of all trials within a given session. We then compared the microsaccade rate curves for the two groups of trials. We provided descriptive statistics and visualizations for both types of analyses.

## Results

Our goal was to document the effects of task-irrelevant sound pulses on visually-driven microsaccadic modulations in rhesus macaque monkeys. Throughout all trials in all experiments, our monkeys fixed their gaze on a central fixation spot presented over a uniform gray background (Methods). On every regular trial, a visual stimulus appeared, and on trials with sound pulses, the visual onset was accompanied by a brief, suprathreshold tone of a single frequency (Methods). On yet other control trials, only the sound pulse was played.

When it was played, the source of the sound pulse was either spatially-nonlateralized, coming from a pair of speakers placed symmetrically behind the display with each speaker at an equal distance to the right or left of the central fixation spot (Methods), or it was spatially-lateralized, coming from a pair of speakers placed either to the right or left of the central fixation spot (and again slightly behind and below the display; Methods). Note that we did not use exactly matched sound and visual stimulus locations, as might be sometimes aimed for in neurophysiological studies of audio-visual integration in multisensory structures like the superior and inferior colliculi (41, 42, 45, 52–54). Rather, we asked: to what extent could nonlateralized or lateralized sounds interfere with or otherwise modify microsaccadic inhibition?

### Spatially-nonlateralized sound alone barely influences microsaccade rate

Our first result was that sound alone had a surprisingly mild, and almost non-existent, impact on microsaccade rate in our monkeys. This can be seen in Fig. 2. When we played the spatially-nonlateralized sound alone (Fig. 2A, B), there was no apparent short-latency change in the rate of microsaccades occurring after auditory stimulus onset in both monkeys. This was very different from what happened when the visual stimulus appeared alone. In this case, and consistent with the prior literature on microsaccadic inhibition in monkeys (1, 8, 9, 17, 18, 20, 24, 25), there was a robust reduction in microsaccade rate within less than 100 ms from visual stimulus onset (Fig. 2C, D). This robust visually-driven microsaccadic inhibition was also clearly evident when both the visual stimulus onset and the auditory pulse occurred together (Fig. 2E, F). Thus, vision dominated the occurrence of stimulus-induced microsaccadic inhibition in our monkeys. This observation is consistent with previous human investigations of saccadic inhibition, in which the inhibition caused by sound alone was generally weak, absent, or inconsistent across subjects and paradigms (34, 55–58). More importantly, this observation has additional value: it makes the use of sound pulses in monkeys a useful experimental tool for modifying visually-driven microsaccadic inhibition, without itself being a major driver for such inhibition. In what follows, we provide further evidence for this idea, again reinforcing the experimental utility of such spatially-nonlateralized sound pulses in invasive pathway studies of sensory-motor transformations in primate brains (also see Discussion).

### Multisensory stimulation with spatially-nonlateralized sound accelerates visually- or auditory-driven microsaccadic rate inhibition

Closer inspection of Fig. 2E, F should reveal that microsaccadic inhibition started earlier in the case of multisensory stimulation (Fig. 2E, F) than in the case of visual stimulus onsets alone (Fig. 2C, D). To better illustrate this, we converted our measured microsaccade onset times into microsaccade rate curves (Methods). Such rate curves are plotted in Fig. 3, and they illustrate multiple key observations. First, with the visual stimulus onset alone, we observed classic microsaccadic inhibition followed by a rebound in microsaccade rate (Fig. 3A, D; blue curves). Second with the spatially-nonlateralized sound alone, there was barely any microsaccadic inhibition in monkey F (Fig. 3A; gray curve), and very mild inhibition in monkey A (Fig. 3D; gray curve), consistent with Fig. 2. Importantly, multisensory stimulation accelerated whatever inhibition was observed with unisensory stimulation alone. For example, in both monkeys, the onset of microsaccadic inhibition with the paired visual and auditory onsets was earlier than the onset of the mild microsaccadic inhibition with the auditory event alone (black arrows in Fig. 3B, E). Similarly, microsaccadic inhibition was earlier on the multisensory trials than in the visual-only trials (Fig. 3C, F). Thus, we observed evidence of multisensory integration (54, 59–61), in the sense that the results with multisensory stimulation did not merely reflect a superposition of the individual unisensory findings; rather, there was a nonlinear interaction in the timing of microsaccadic inhibition in the case of multisensory stimulation. Moreover, the multisensory result was more similar to the visual-only result, as evidenced by the strong inhibition that was absent with the auditory sound pulses alone. In other words, vision dominated sound in the cue-driven microsaccadic inhibition effect.

**Figure 3.**
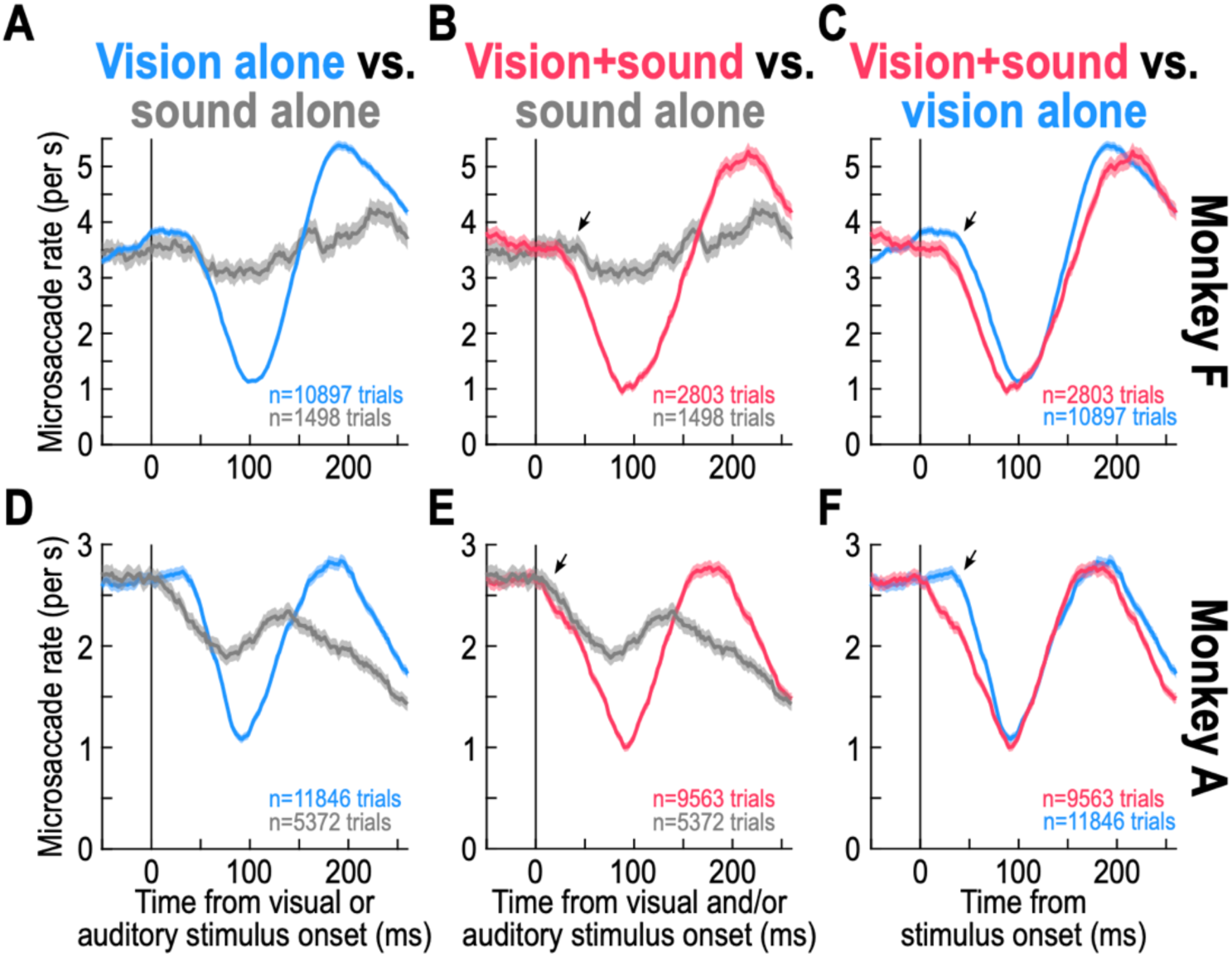
Multisensory stimulation accelerates the timing of unisensory-driven microsaccadic inhibition, with vision dominating the strength of the inhibition. **(A)** In monkey F, we estimated microsaccade rate from individual trials like those shown in Fig. 2 (Methods). Each curve shows average microsaccade rate across all trials of a given type (blue for visual stimulus alone, and gray for spatially-nonlateralized sound alone); error bars indicate across-trial SEM. In the visual modality, classic microsaccadic inhibition, followed by a rate rebound, was observed (note that monkeys generally show higher microsaccade rates, and faster rate modulation dynamics, than humans). However, with the sound alone, there was only a very mild reduction in microsaccade rate, at approximately the same time as the visually-driven inhibition. **(B)** When we paired the visual stimulus with a simultaneous sound pulse (Methods), the microsaccadic rate signature (red) looked very similar to the visually-driven signature in **A**. However, the inhibition was clearly accelerated relative to the small rate modulation seen in the case of the sound pulse alone (small black arrow). **(C)** Consistent with this, plotting the rate signatures for vision in isolation (blue) and multisensory stimulation (red) together revealed a clear acceleration of microsaccadic inhibition with multisensory stimulation. Thus, multisensory stimulation does not cause a superposition of effects expected from each sensory modality individually, but it accelerates microsaccadic inhibition. **(D-F)** Similar results for monkey A. Note how the mild sound-only inhibition in this monkey started earlier than in monkey F (**D**, **E**). Despite that, multisensory stimulation still accelerated the inhibition relative to sound alone (**E**; compare red and gray curves) or relative to vision alone (**F**; compare red and blue curves). The numbers of trials in each condition are indicated in each panel.

To quantitatively document the acceleration of visually-driven microsaccadic inhibition timing by spatially-nonlateralized sound pulses, we calculated a latency measure of microsaccadic inhibition (L_B25_). This measure indicates the time at which the microsaccade rate after stimulus onset dropped by 25% relative to its prestimulus level (Methods). For monkey F, this latency was 68 ms in the visual-only condition, but it was 50 ms in the multisensory condition (L_B25_ difference = 18 ms, *p* < 0.0001, 95% CI [-6,8]). Similarly, in monkey A, L_B25_ was 62 ms in the visual-only condition and 48 ms in the case with a paired spatially-nonlateralized sound (L_B25_ difference = 14 ms, *p* < 0.0001, 95% CI [-6,6]). Thus, the spatially-nonlateralized sound accelerated the visually-driven microsaccadic inhibition.

Interestingly, even though we could not reliably estimate the L_B25_ parameter for the sound-only condition of our task (due to the very weak inhibition seen in Figs. 2, 3, which never reduced microsaccade rate by more than our measurement threshold of 25%; Methods), visual inspection suggested that multisensory integration additionally also accelerated the timing of microsaccadic inhibition relative to the sound-only trials. Specifically, while there was no apparent latency difference between sound-only trials and visual-only trials for monkey F (Fig. 3A), the multisensory trials had a shorter latency of inhibition than the sound-only trials for this monkey (Fig. 3B). In monkey A’s case, the sound-only trials exhibited earlier inhibition than the visual-only trials (Fig. 3D), and the multisensory trials still showed a hint of a latency benefit compared to the sound-only trials (Fig. 3E), in agreement with monkey F’s results. Thus, the spatially-nonlateralized sound also accelerated microsaccadic inhibition relative to sound-only conditions.

Finally, and as mentioned above, the strength of microsaccadic inhibition was vastly different between the sound-only trials and the multisensory trials. In the latter trials, maximal inhibition (i.e. the minimum value reached by the microsaccade rate curve shortly after stimulus onset, relative to the prestimulus rate value) was similar to the maximal inhibition observed in the visual-only trials, suggesting that the visual modality dominated the stimulus-driven modulations in microsaccade rate that we observed. We statistically confirmed this observation. The difference in maximal inhibition between auditory-only and visual-only trials was 1.8831 microsaccades/s in monkey F (*p_BH_* = 0.0003, 95% CI [-0.2959,0.1997]) and 0.8016 microsaccades/s in monkey A (*p_BH_* = 0.0003, 95% CI [-0.1663,0.1491]). Similarly, between the auditory-only and multisensory trials, the differences were 2.2345 microsaccades/s (*p_BH_* = 0.0003, 95% CI [-0.4798,0.4240]) in monkey F and 0.8821 microsaccades/s (*p_BH_* = 0.0003, 95% CI [-0.1668,0.1620]) in monkey A. In contrast, the differences in maximal inhibition between the visual-only and multisensory trials were tiny and non-significant in both monkeys: 0.1853 microsaccades/s (*p_BH_* = 0.0526, 95% CI [-0.1731,0.1989]) in monkey F, and 0.0804 microsaccades/s (*p_BH_* = 0.1725, 95% CI [-0.1146,0.1178]) in monkey A. Importantly, these differences between the visual-only and multisensory trials were markedly smaller than those expected from the maximal inhibition of the auditory-only trials; in those trials, maximal inhibition relative to prestimulus baselines (i.e. the absolute value between prestimulus and minimum microsaccade rate) was 0.3571 microsaccades/s in monkey F and 0.7772 microsaccades/s in monkey A. Thus, the multisensory effects were not a superposition of the individual unisensory ones. This could be because our visual and auditory stimuli were not spatially aligned (Methods) like in other studies (55), and also because we did not use low-contrast visual stimuli and/or low-intensity sound pulses, which are known to amplify multisensory integration effects (54, 55, 62, 63). It is also interesting to note here that we saw no clear differences in the post-inhibition microsaccadic rate rebound between visual-only and multisensory trials (Fig. 3C, F; the first microsaccade rate peak occurring after the inhibition phase).

### Microsaccadic rate inhibition does not adapt to repeated visual or auditory sensory stimulation

Because of the surprisingly mild sound-only rate modulations that we saw in our data so far, we next checked whether this could reflect adaptation to repeated exposure to our sound pulses. This question touches on a broader one about whether saccadic inhibition, in general, exhibits sensory adaptation or not. In the visual modality, this does not seem to be the case, given how saccadic inhibition still occurs in monkeys after thousands of trial repetitions of the same kind (8). However, we explicitly tested this possibility here, and also for the sound stimuli.

In a first analysis, we explored whether exposure to sound on a previous trial was sufficient to weaken microsaccadic inhibition for sound pulses in sound-only trials. For every sound-only trial, we checked whether it was preceded by either a trial with only a visual stimulus or a trial with a sound (either the sound alone or the sound with a visual stimulus; Methods). We combined sound alone and vision plus sound in the latter group because of the scarcity of our sound-only trials (Methods) and also to maximize the likelihood of adaptation to a prior exposure to a sound. For both monkeys, the rate curves in the two trial groupings were unaltered during the early stimulus-driven microsaccadic inhibition phase (Fig. 4A, E). If anything, monkey A might have even shown trends towards sensitization, rather than adaptation. Thus, there was no evidence for adaptation in the sound-only trials. Interestingly, at later times from stimulus onset, monkey A did show changes in microsaccade rates when the previous trial contained a sound; these later epochs of the microsaccadic rate signature are driven by top-down control (15) and might more strongly reflect adaptation than the early reflexive inhibition phase.

**Figure 4.**
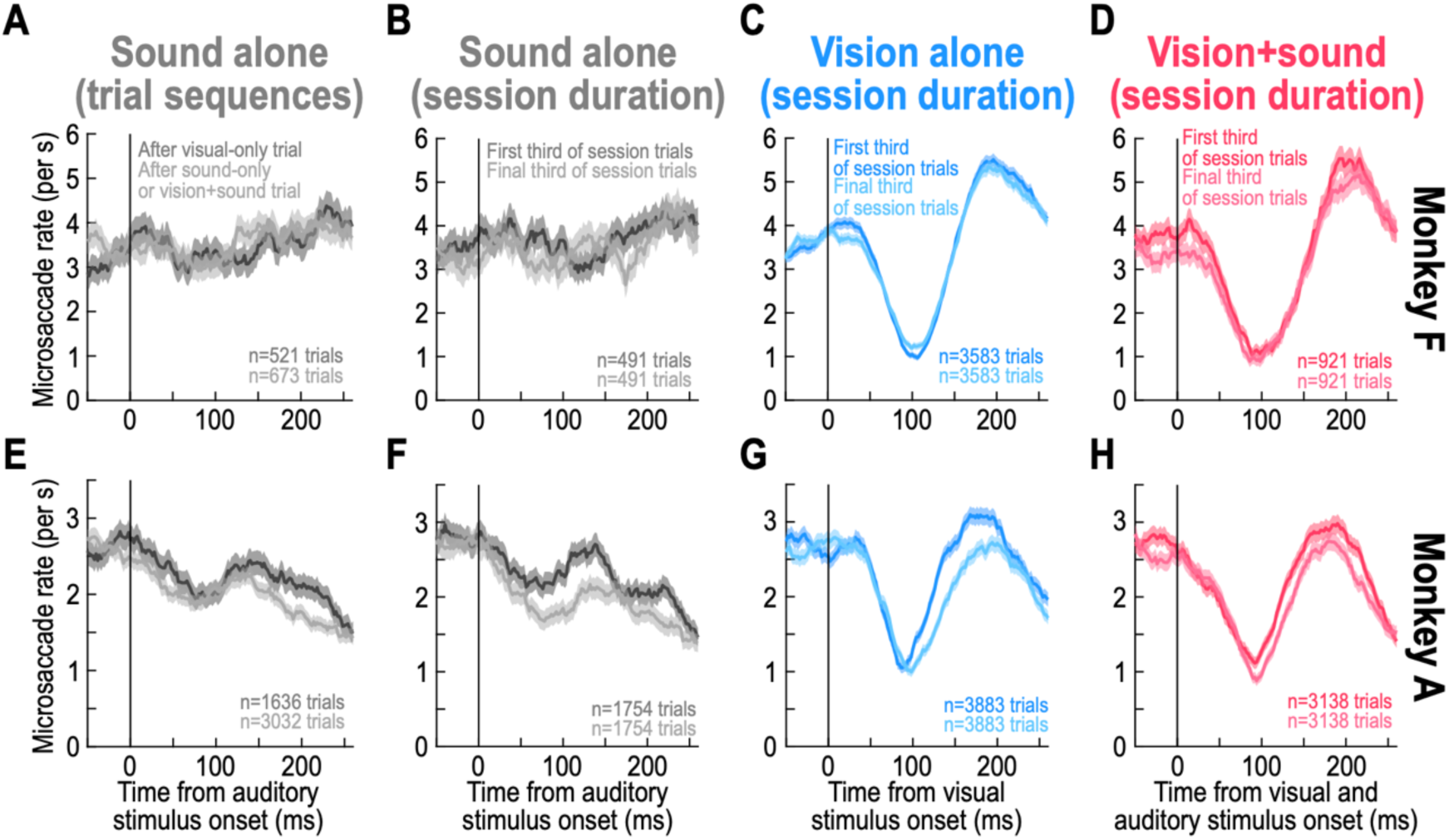
Lack of adaptation in microsaccadic inhibition, whether with visual or auditory stimuli. **(A)** For monkey F, we split all sound-only trials as a function of whether they were preceded by a visual-only trial (dark) or a sound trial (bright; Methods). There was no difference in microsaccade rate across the two groups, suggesting that prior exposure to sounds did not weaken the sound-only effects seen in Fig. 3. **(B)** For the same monkey, we also checked for adaptation by grouping trials within a session as being either early (dark) or late (bright) within a session. Again, the sound-only rates were unaltered by session duration, suggesting a lack of adaptation to repeated exposure to sound stimuli. **(C)** Similar to **B**, but for the visual-only trials. Again, there was no adaptation in either the timing or strength of microsaccadic inhibition to repeated exposure to visual stimuli. **(D)** This was also the case for the multisensory trials. **(E-H)** Similar to **A**-**D**, but for monkey A. If anything, monkey A might have been sensitized, rather than adapted, in the sound only trials (trends for earlier inhibition after prior sound exposure, whether in the trial-based or session-based analyses of **E**, **F**). This monkey also showed weaker rebounds after the inhibition phase in the later trials, suggesting adaptation in top-down (15) recovery from microsaccadic inhibition. Error bars denote SEM across trials. All other conventions are like in Fig. 3.

We also considered longer-term adaptation effects, which might reflect longer-term alterations in cognitive state after repeated exposure to the same set of trial types and stimuli within a given session. To do so, we asked whether trials late in a session had different microsaccadic inhibition profiles than trials early in a session (Methods). We picked either the first third or final third of trials from any given session, and we plotted microsaccade rates for the two groups of trials. The results for monkey F are shown in Fig. 4B-D, and those for monkey A are shown in Fig. 4F-H. Again, there was no evidence of sensory adaptation in the early microsaccadic inhibition phase, whether for sound-only trials, vision-only trials, or multisensory trials. In fact, for the sound-only trials, monkey A might have been sensitized, rather than adapted, by repeated exposure to the same trial types throughout a given session. Moreover, this monkey showed weaker and delayed rebound phases in the later trials, again suggesting adaptation in the top-down components of the microsaccadic rate signature (15).

Therefore, our results indicate that short-latency, reflexive microsaccadic inhibition does not adapt to repeated sensory exposure, in both the visual and auditory modalities.

### Spatially-nonlateralized sound modulates, but does not eliminate, visually-driven microsaccade direction modulations

Our next goal was to evaluate microsaccade direction modulations. Prior research in humans (19, 64–68) and monkeys (8, 9, 16–18) has revealed the occurrence of consistent modulations in microsaccade directions after extrafoveal visual stimulus onsets. If vision really did dominate the cue-driven effects of multisensory stimulation on microsaccades in our monkeys, then analyzing the monkeys’ microsaccade directions in the experiments of Figs. 2, 3 should still reveal such modulations, despite the spatially-nonlateralized sound pulses. This is what we found.

With visual stimulation alone, we replicated the previous classic results. For example, for monkey F, the visually-driven inhibition of microsaccades congruent to the visual stimulus hemifield (Fig. 5A; congruent) was delayed relative to the inhibition of microsaccades towards the opposite hemifield (Fig. 5A; incongruent). Mechanistically, this means that microsaccade plans in the direction of the appearing visual stimulus were more likely to survive the countermanding effect of the stimulus onset than microsaccade plans in the opposite direction (18); across trials, the net result is a biasing of microsaccade directions towards the appearing visual stimulus shortly after its onset. This biasing can also be seen for the same monkey in Fig. 5C (blue curve). In this analysis, we plotted the likelihood for an occurring microsaccade within a given time bin to be in the direction of the visual stimulus hemifield (Methods). During the short-latency period of microsaccadic rate inhibition, almost all microsaccades that did occur in this monkey were congruent with the visual stimulus hemifield (black arrow), and later microsaccades were in the opposite direction (Fig. 5C; blue curve). Similar effects were also observed in monkey A (Fig. 5D and Fig. 5F; blue curve), even though this monkey had weaker cueing effects than monkey F.

**Figure 5.**
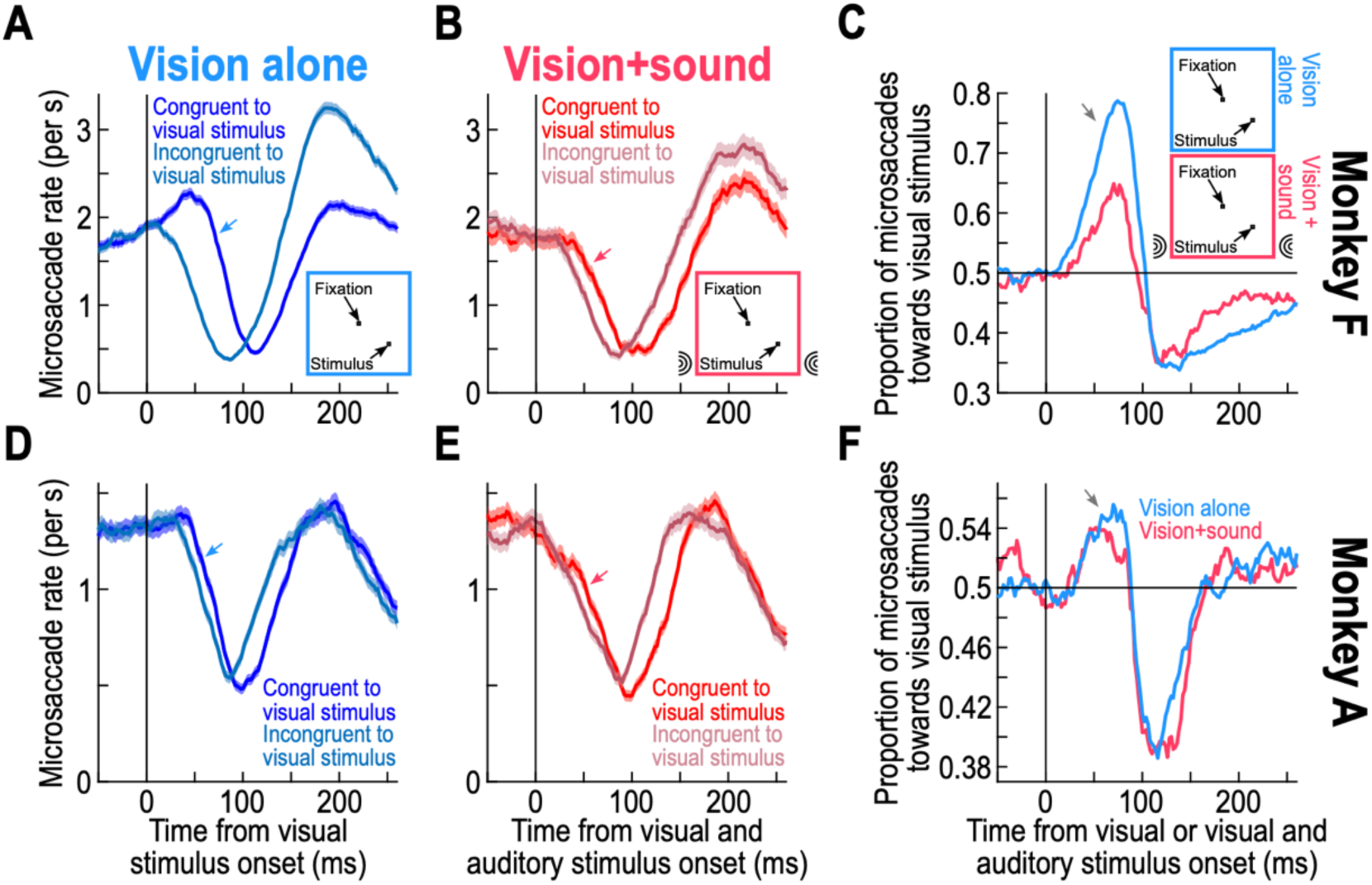
Spatially-nonlateralized sound modulates, but does not abolish, visually-driven microsaccade direction modulations in monkeys. **(A)** With only visual onsets, the inhibition of microsaccades congruent to the hemifield of the appearing visual stimulus was later than the inhibition for opposite microsaccades (blue arrow). This means that near the time of microsaccadic inhibition, the microsaccades that did occur were more likely to be in the direction of the appearing visual stimulus (9, 17, 18). **(B)** When adding a simultaneous spatially-nonlateralized sound with the visual onset, this effect remained, although it was markedly weakened. Note that in **A**, **B**, the prestimulus microsaccade rates are lower than those in Fig. 3; this is because we now characterized the rates for only subsets of microsaccades in any given condition (based on their direction classifications), whereas Fig. 3 lumped all microsaccade directions together. **(C)** For the same monkey, we plotted on the x-axis the time from visual stimulus onset, and on the y-axis the proportion of all occurring microsaccades within a given time bin that were directed towards the visual field of the appearing stimulus (Methods). We confirmed the results of **A**, **B**: at the time of microsaccadic inhibition, most microsaccades that did occur were directed towards the appearing visual stimulus, and the spatially-nonlateralized sound interfered with, but did not completely eliminate, this phenomenon. **(D-F)** Similar observations in monkey A. Note that this monkey had a weaker cueing effect on microsaccade directions (**D**), but this effect was still visible with the interfering sound (**E**, **F**). The generally weaker effect was explained by stronger prestimulus biases in microsaccade directions in this monkey than in monkey F (see Fig. 6). Each curve in **A**, **B**, **D**, **E** is the mean across trials, and the error bars indicate SEM. The total number of trials in each panel was the same as in Fig. 3.

With the spatially-nonlateralized sound pulses, the visually-driven microsaccade direction modulations persisted, but they were weakened. For example, in monkey F, L_B25_ for congruent microsaccades with vision alone was 84 ms, and it was 46 ms for incongruent microsaccades (Fig. 5A); giving a time difference of 38 ms (*p* < 0.0001, 95% CI [-4,4]). With the spatially-nonlateralized sound pulse occurring in synchrony with the visual stimulus onset, the L_B25_ values were 62 ms and 40 ms for the congruent and incongruent microsaccades, respectively; giving a time difference of only 22 ms (*p* = 0.0016, 95% CI [-12,12]) (Fig. 5B). For monkey A, the L_B25_ difference was 10 ms with vision alone (68 ms and 58 ms for congruent and incongruent microsaccades, respectively; *p* = 0.046, 95% CI [-8,8]; Fig. 5D), and it was 8 ms for vision plus the spatially-nonlateralized sound pulses (52 ms and 44 ms for congruent and incongruent microsaccades, respectively; *p* = 0.2651, 95% CI [-12,12]; Fig. 5E). These effects were also evident in the proportion plots of Fig. 5C, F (red curves). As can be seen, in both monkeys, the cue-driven direction modulations of microsaccades reflected the visual landscape of sensory stimulation, but they were weaker with multisensory stimuli than with visual onsets alone.

Thus, the spatially-nonlateralized sounds in our experiments interfered with, but did not completely eliminate, the visually-driven microsaccade direction modulations shortly after stimulus onset. This indicates that spatially-nonlateralized sounds can have multiple divergent effects on reflexive cue-driven oculomotor behavior: on the one hand, they can enhance the salience of the sensory transients (say, via increased alertness) and accelerate the timing of microsaccadic inhibition (Fig. 3); on the other hand, their spatially-diffuse nature can increase the noise in the spatial location representation induced by the visual stimulus onset (putatively in spatially-organized visual-motor maps of the brain, like the superior colliculus; Fig. 5) (42, 53, 54, 59, 69).

The results of Fig. 5D-F for monkey A are particularly convincing that visual cueing effects, despite being initially weak in this monkey, still showed the same trends with the spatially-nonlateralized sound pulses. However, we further analyzed the data from this and the other monkey, in order to better understand this weaker visual-only cueing effect in this particular animal. We believe that this reason can become especially important in neurophysiological experiments involving lateralized brain structures, like the primary visual cortex or the superior colliculus. For example, in causal perturbation neurophysiological experiments, one might experimentally activate or inactivate only one side of the primary visual cortex or superior colliculus, requiring analysis of trials with visual stimuli in only one hemifield (rather than by combining hemifields like we did in Fig. 5). Thus, we asked whether the generally weaker cueing effects for monkey A (with or without the spatially-nonlateralized sounds) could have been related to potential hemifield asymmetries in this monkey’s microsaccades.

We found that prestimulus microsaccade directions could indeed exhibit idiosyncratic biases in individual monkeys, like in humans (70), and which likely reflect biases in ocular drift directions (10, 11). To demonstrate this, we split trials according to the visual field location of the visual stimulus, and we also split the observed microsaccades within each group of trials according to whether they were rightward or leftward in direction. Monkey F tended to make more rightward than leftward microsaccades before stimulus onset (Fig. 6A, B; vertical bidirectional black arrows). In contrast, monkey A had an even stronger bias, but in the opposite direction: there were almost twice as many leftward than rightward microsaccades in this monkey before stimulus onset (Fig. 6D, E; vertical bidirectional black arrows), and the prestimulus rightward microsaccade rate was uncharacteristically low for macaque monkeys. Thus, for any given monkey, a leftward or rightward visual stimulus onset could be in or out of line with the monkey’s intrinsic microsaccade direction bias. The next question became, then, whether this could cancel visually-driven cueing effects within specific hemifields or not.

**Figure 6.**
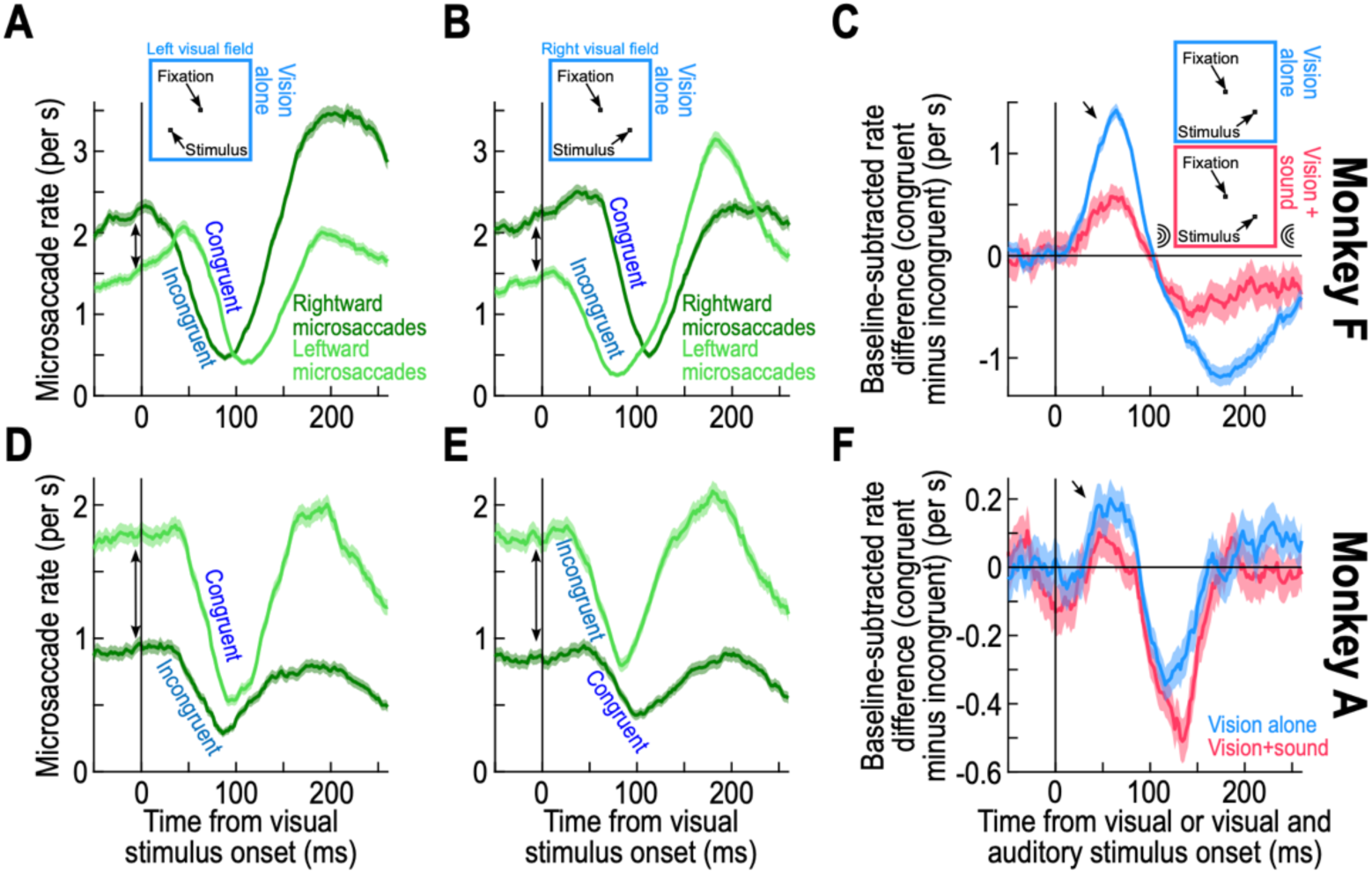
Persistence of the visual dominance over microsaccade direction modulations, even in the face of potentially strong prestimulus biases in microsaccade directions. **(A, B)** Similar data to those in Fig. 5A (visual stimulus onset alone), but after separating trials based on the visual field location of the appearing stimulus: left visual field in **A** and right visual field in **B**. In the former case, leftward microsaccades were congruent with the stimulus location (labeled “congruent”), and were inhibited later than rightward microsaccades (**A**); in the latter case (right visual field stimulus), the later inhibition was for the rightward movements (**B**), consistent with Fig. 5. This way of plotting the data was particularly useful for revealing prestimulus microsaccade direction biases (more leftward movements in this monkey). **(C)** A measure of stimulus-induced direction oscillations (like in Fig. 5C), but that can be applied in each hemifield individually despite prestimulus biases. In this measure, we calculated the baseline-subtracted difference in microsaccade rates between different microsaccade directions (here, between congruent and incongruent microsaccades across all visual stimulus locations). Note that we applied this measure for both hemifields combined in this case, for easier comparison to Fig. 5C; however, the measure is equally useful within each hemifield individually. **(D-F)** Similar results for monkey A. There was an even larger prestimulus bias in microsaccade directions (**D**, **E**; approximately twice as many leftward than rightward movements). Despite that, there was still later inhibition for the rate curve of the congruent microsaccades to the visual stimulus (leftward in **D** and rightward in **E**). Consistent with this (and Fig. 5F), the baseline-subtracted rate difference between congruent and incongruent microsaccades (**F**) shifted towards the positive direction during microsaccadic inhibition, even with the spatially-nonlateralized sound pulses (note that the y-axis range is smaller than that in **C**, reflecting this monkey’s weaker cueing effects in general). Error bars denote SEM. All other conventions are similar to earlier figures.

The individual-hemifield cueing effects on microsaccade directions remained consistent with the global results of Fig. 5, despite individual monkey differences in prestimulus microsaccade direction biases. In monkey F, when the visual stimulus onset was in the left visual field (Fig. 6A), leftward microsaccades were inhibited later than rightward microsaccades (L_B25_ of 86 ms versus 46 ms, respectively; L_B25_ difference = 40 ms, *p* < 0.0001, 95% CI [-8,8]), even though leftward microsaccades were less frequent than rightward microsaccade before stimulus onset in this monkey. Similarly, in the same monkey, when the visual stimulus onset was in the right visual field (Fig. 6B), now the rightward microsaccades were inhibited later than the leftward movements (L_B25_ of 82 ms versus 44 ms, respectively; L_B25_ difference = 38 ms, *p* < 0.0001, 95% CI [-8,6]). In monkey A, with the much bigger prestimulus bias in microsaccade directions, the results were similar. For the left visual field stimulus (Fig. 6D), L_B25_ was 62 ms versus 54 ms for leftward and rightward microsaccades, respectively (L_B25_ difference = 8 ms, *p* = 0.1232, 95% CI [-10,10]); note that the result of the significance test could reflect the different dynamics of rate inhibition between the two curves. For the right visual field stimulus (Fig. 6E), the results were 78 ms versus 58 ms for rightward and leftward microsaccades, respectively (L_B25_ difference = 20 ms, *p* < 0.0009, 95% CI [-12,12]). Thus, even when analyzing individual hemifields, as might be the case in neurophysiological experiments in lateralized brain structures like the visual cortex or the superior colliculus, visually-driven cueing effects will still be observed despite sometimes strong prestimulus microsaccade biases; congruent microsaccade rate (with the visual stimulus location) is inhibited later than incongruent microsaccade rate, and for each visual stimulus hemifield individually (Fig. 6A, B, D, E).

The above results, besides supporting the conclusions from Fig. 5, allowed us to next devise a measure of cue-driven microsaccade direction oscillations that is independent of prestimulus direction biases (Methods). With this metric, we first measured the rate of congruent microsaccades and the rate of incongruent microsaccades as a function of time from stimulus onset. Then, before comparing these two rates to identify cueing effects on microsaccade directions, we subtracted the prestimulus microsaccade rate from each curve individually. Only after such baseline subtraction did we calculate the difference in microsaccade rates between congruent and incongruent movements. As seen in Fig. 6C, F for both monkeys, the difference metric was close to zero in the prestimulus interval and thus isolated the cue-driven modulatory direction effects despite idiosyncratic prestimulus biases. The results from both monkeys (Fig. 6C, F) were consistent with those from all of our other analyses above. Specifically, for the vision-only trials (blue curves), there was a biphasic oscillation in microsaccade directions, first towards and then opposite the recently appearing stimuli. And, with the visual onsets being paired with spatially-nonlateralized sound pulses, the direction oscillations persisted but were weaker.

### Spatially-lateralized sound similarly influences microsaccade rate as spatially-nonlateralized sound, with and without visual stimulation

Our results so far indicate that spatially-nonlateralized sound pulses alone had very mild impacts on stimulus-driven microsaccade rate modulations, but that they still accelerated visually-driven microsaccadic inhibition. Moreover, such sound pulses interfered with, but did not completely eliminate, visually-driven microsaccade direction modulations. We interpret these results as suggesting that vision dominates sound in transient cue-induced microsaccadic eye movement modulations in rhesus macaque monkeys. Next, we show that this conclusion holds even when using spatially-lateralized sound pulses that were either congruent or incongruent with the hemifield location of the appearing visual stimuli.

In separate blocks of trials, we repeated the same experiments as above, but with the sound pulse on every trial coming from either the right or left side of the display placed in front of the animals (Methods; Fig. 1). Again, in the absence of a visual onset, such spatially-lateralized sound pulses barely affected ongoing microsaccade rates in both monkeys. This can be seen in Fig. 7A, B, in which no microsaccadic inhibition occurred (also see Fig. 9C, F later for a more quantitative microsaccade rate version of the same data). In contrast, trials with a visual onset paired with a spatially-lateralized sound pulse showed clear microsaccadic inhibition, regardless of whether the sound source was in a hemifield congruent (Fig. 7C, D) or incongruent (Fig. 7E, F) with the hemifield of the appearing visual stimulus. Thus, we replicated the same results of Fig. 2, but now with a spatially-lateralized sound source.

**Figure 7.**
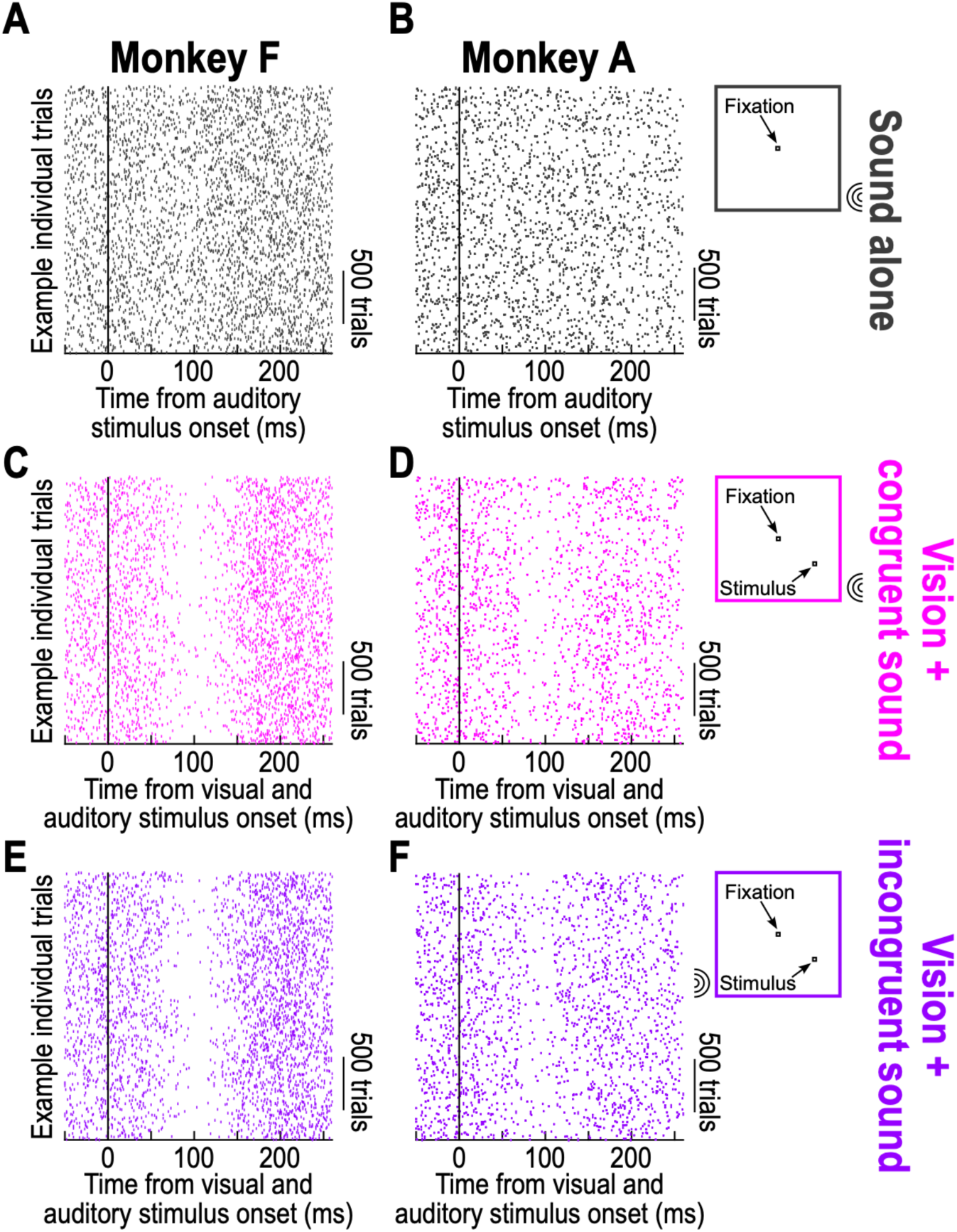
Spatially-lateralized sound alone again does not cause clear microsaccadic inhibition in monkeys. **(A, B)** Similar to Fig. 2A, B but this time when the sound source was placed on only one side (either right or left) of the visual display. In the shown trials, the sound pulse occurred alone, and there were no visual transients on the display. There were again barely any changes in microsaccade rate caused by the sound pulse alone (also see Fig. 9). **(C, D)** Microsaccadic inhibition was robust when a visual stimulus onset was paired with the spatially-lateralized sound pulse. In this case, the visual stimulus appeared in the same hemifield as the spatially-lateralized sound pulse (Methods). **(E, F)** Microsaccadic inhibition was also robust when the visual onset was in the opposite hemifield from the spatially-lateralized sound pulse. All other conventions similar to Fig. 2.

We also replicated the observation that multisensory stimulation accelerated the onset of visually-driven microsaccadic inhibition in both monkeys. Consider, for example, Fig. 8A for monkey F and Fig. 8C for monkey A. Here, we plotted microsaccade rates in the visual-only condition in blue (these data are the same as those in Fig. 3 for the same condition). We also plotted microsaccade rates when the visual onset was paired with a spatially-lateralized sound pulse that was either congruent (magenta) or incongruent (purple) with the hemifield of the appearing visual stimulus. In both cases, microsaccadic inhibition occurred earlier than with a visual stimulus alone. Specifically, for monkey F (Fig. 8A), L_B25_ for the visual-only condition was 68 ms, whereas it was 58 ms (L_B25_ difference = 10 ms, *p_BH_* = 0.0356, 95% CI [-6,8]) and 56 ms (L_B25_ difference = 12 ms, *p_BH_* = 0.0050, 95% CI [-6,6]) for the congruent and incongruent sound trials, respectively. Similarly, for monkey A (Fig. 8C), L_B25_ for the visual-only condition was 62 ms, whereas it was 50 ms (L_B25_ difference = 12 ms, *p_BH_* = 0.0070, 95% CI [-6,6]) and 50 ms (L_B25_ difference = 12 ms, *p_BH_* = 0.0072, 95% CI [-6,6]) for the congruent and incongruent sound trials, respectively. Thus, sound, whether spatially-nonlateralized (Fig. 3) or spatially-lateralized (Fig. 8), accelerated visually-driven microsaccadic inhibition in our macaque monkeys.

**Figure 8.**
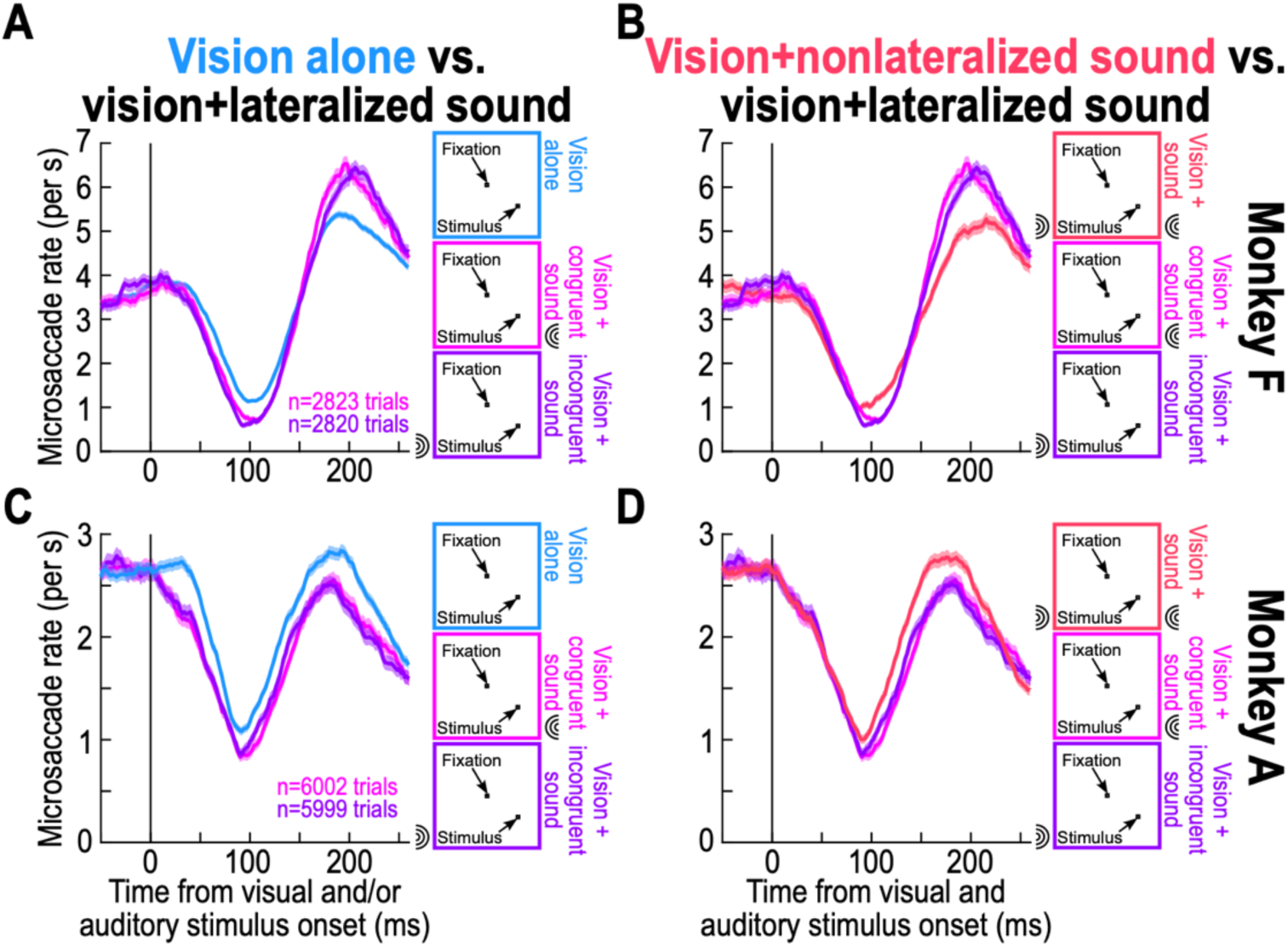
Spatially-lateralized sound accelerates the timing of visually-driven microsaccadic inhibition similarly to spatially-nonlateralized sound. **(A)** In blue, we plotted the microsaccade rate of monkey F on the trials with only a visual onset (same data as in Fig. 3A). In magenta and purple, we plotted microsaccade rate when the visual onset was paired with a spatially-lateralized sound (congruent with the visual stimulus hemifield in magenta, and incongruent in dark purple). The rate inhibition occurred earlier with multisensory than visual-only stimulation, consistent with the case of a spatially-nonlateralized sound (Fig. 3). **(B)** The acceleration of the timing of microsaccade rate inhibition by sound was the same whether the sound was spatially-lateralized or nonlateralized. The magenta and dark purple curves are the same as those in **A**, and the red curve shows microsaccade rate in monkey F when the visual stimulus onset was paired with a spatially-nonlateralized sound (same data as in Fig. 3B). Microsaccadic inhibition timing was indistinguishable across all three conditions **(C, D)** The same conclusions were reached for monkey A. Error bars denote SEM, and the numbers of trials for the blue and red curves are the same as those reported earlier in Fig. 3.

The acceleration by sound of visually-driven microsaccadic inhibition (Figs. 3, 8) was also independent of the spatial biases in the sound pulses themselves. Specifically, when we plotted microsaccade rates from the vision plus spatially-nonlateralized sound trials (like in Fig. 3) along with the vision plus spatially-lateralized sound trials, we found almost perfect overlap in the timing of microsaccadic inhibition in both monkeys (Fig. 8B, D). Quantitatively, for monkey F, the difference between L_B25_ in the vision plus spatially-nonlateralized sound trials and L_B25_ in the congruent sound trials was -8 ms (*p_BH_* = 0.1932, 95% CI [-8,8]), and it was -6 ms (*p_BH_* = 0.3972, 95% CI [-8,8]) between the vision plus spatially-nonlateralized sound trials and the incongruent sound trials. Similarly, no differences were observed for monkey A (L_B25_ difference between nonlateralized vs congruent sound = -2 ms, *p_BH_* = 1, 95% CI [-8,10]; L_B25_ difference between nonlateralized vs incongruent sound = -2 ms, *p_BH_* = 1, 95% CI [-8,8]). Moreover, even in the vision plus spatially-lateralized sound condition itself, the latency of microsaccadic inhibition did not differ between the trials in which the sound pulse was congruent or incongruent with the visual stimulus location (for monkey F, L_B25_ difference = 2 ms, *p_BH_* = 0.7226, 95% CI [-6,6]; for monkey A, L_B25_ difference = -2 ms, *p_BH_* = 1, 95% CI [-8,8]). Thus, with suprathreshold sounds, it did not matter whether they were spatially lateralized or spatially nonlateralized for their impact on the timing of visually-driven microsaccadic inhibition.

In terms of the strength of microsaccadic inhibition, it was again dominated by the visual condition and not by the sound direction. However, we did notice that maximal inhibition was slightly stronger in the spatially-lateralized than spatially-nonlateralized trials (Fig. 8B, D). For monkey F, the difference between the maximal inhibition in the spatially-nonlateralized trials and the maximal inhibition in the congruent sound trials was 0.2405 microsaccades/s (*p_BH_* = 0.0522, 95% CI [-0.2071,0.2123]); the difference between the maximal inhibition in the spatially-nonlateralized trials and the maximal inhibition in the incongruent sound trials was 0.3603 microsaccades/s (*p_BH_* = 0.0012, 95% CI [-0.1944,0.1969]). For monkey A, these values were 0.1687 microsaccades/s (*p_BH_* = 0.0898, 95% CI [-0.1313,0.1320]) and 0.1383 microsaccades/s (*p_BH_* = 0.0012, 95% CI [-0.1328,0.1366]), respectively. Though some p-values did not withstand correction for multiple comparisons, the trend was consistent across the monkeys. This is unlikely to be due to the small differences in sound intensity that existed between the experiments (Methods), because these differences could be of opposite signs across the two monkeys (Methods) despite both monkeys showing the same effect of stronger inhibition for the spatially-lateralized sound configuration.

The rebound in microsaccade rate (after microsaccadic inhibition) was also different, but monkey-specific (Fig. 8B, D). As we discuss in more detail below, this latter effect might reflect the individual monkey’s rebound in microsaccade rate after sound-only trials, as well as other top-down strategies in recovering from microsaccadic inhibition (9, 15, 17, 27, 71). For example, monkey F had a slight increase in microsaccade rate approximately 100-200 ms after spatially-lateralized sound onset alone (can be seen in Fig. 9C below), whereas monkey A had a slight decrease at such a time (can be seen in Fig. 9F below). Thus, irrespective of spatial bias in the sound pulses, multisensory stimulation resulted in microsaccade rate inhibition that was largely dominated by the inhibition caused by visual stimulation alone; any small differences in the timing or strength of maximal inhibition did not reflect a superposition of unisensory effects, but the sound acted to modulate the properties of the visually-driven outcomes.

**Figure 9.**
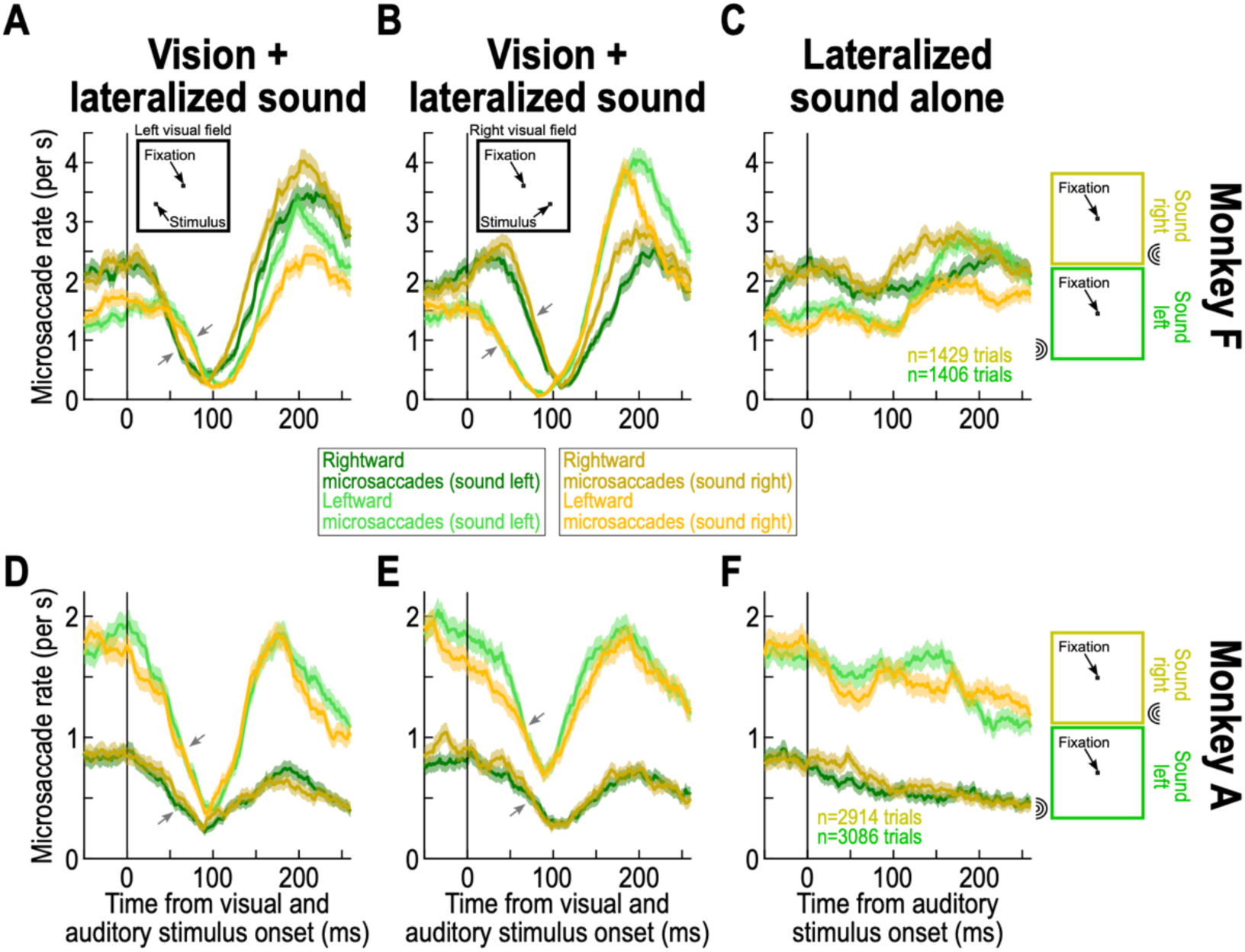
Spatially-lateralized sound location does not matter for the impacts of visual stimulus location on microsaccade direction modulations during microsaccadic inhibition. **(A)** Similar to Fig. 6A, but now exploring the effect of the hemifield location of the spatially-lateralized sound. For a left visual field stimulus, green shows microsaccade rates for rightward or leftward microsaccades and a left sound source; yellow shows the same curves but now with a right sound source. At the time of microsaccadic inhibition, it did not matter whether the sound was on the right or left for each curve. That is, the congruency to the visual stimulus location was what dictated the timing of rate inhibition (black oblique arrows highlight the similarity of inhibition timing between corresponding pairs of yellowish and greenish curves). **(B)** Similarly, when the visual stimulus was in the right hemifield, microsaccadic inhibition reflected the congruency of the microsaccades with the visual stimulus, regardless of whether the sound was congruent (yellow) or incongruent (green) with the visual stimulus location (black oblique arrows). **(C)** With a spatially-lateralized sound alone, this monkey did not show any appreciable microsaccadic inhibition, regardless of sound or movement direction, consistent with Fig. 7. Critically, the green and yellow curves were overlapping at the time of expected microsaccadic inhibition in **A**, **B**. At later times, rate rebounds could depend on sound location, explaining some of the differentials between the green and yellow curves in the post-inhibition rebound phases in **A**, **B**. **(D-F)** Similar results for monkey A. This monkey was more sensitive to rightward sounds (slightly inhibited leftward microsaccades with the rightward sound alone in **F**). This caused a slightly earlier onset of inhibition in the yellow curves of **D**, **E**, but the overall timing of visually-driven microsaccadic inhibition was unaffected by sound location (oblique black arrows). Error bars denote SEM, and the total numbers of trials for the vision+spatially-lateralized sound trials were the same as those in Fig. 8. Also see Fig. 10 for an alternative visualization of **A**, **B**, **D**, **E**.

### Visually-driven microsaccade direction modulations do not change with the hemifield congruency of spatially-lateralized sound

The strongest evidence that we had for the dominance of vision over sound in cue-driven microsaccadic modulations in rhesus macaque monkeys emerged when we looked at the microsaccade direction modulations shortly after stimulus onset. We repeated the analyses of Fig. 6A, B, D, E, but this time for multisensory stimulation with either a spatially-lateralized sound pulse in the right hemifield or a spatially-lateralized sound pulse in the left hemifield. From the earlier analyses (Fig. 6A, B, D, E), we already established that there was a visual cueing effect despite prestimulus biases in microsaccade directions in the individual monkeys (which were particularly strong in monkey A). Now, we found that the visual cueing effect in each hemifield of each monkey (that is, the timing of microsaccadic inhibition for microsaccades congruent and incongruent with visual stimulus location) did not depend on the location of the spatially-lateralized sound during the critical microsaccadic inhibition periods. This conclusion can be reached from the data shown in Fig. 9A, B for monkey F and those shown in Fig. 9D, E for monkey A (Fig. 10 also shows an alternate visualization aid of the effects). In these data, we plotted the microsaccade rate curves exactly as we did earlier in Fig. 6. When the visual stimulus was in the left hemifield (Fig. 9A, D), microsaccadic inhibition was later for leftward than rightward microsaccades whether the paired sound pulse with stimulus onset was in the right (yellowish curves) or the left (greenish curves) hemifield; in each panel, the corresponding yellow and green curves had almost perfect overlap during the microsaccadic inhibition phase (black oblique arrows). Similarly, when the visual stimulus was on the right (Fig. 9B, E), microsaccadic inhibition was later for rightward than leftward microsaccades (the expected visually-driven cueing effect), and the microsaccadic inhibition timing for each of the microsaccade direction curves was almost identical regardless of whether the sound pulse was in the right or left hemifield (compare the yellowish and greenish curves in each panel; black oblique arrows). Therefore, the transient cue-driven modulation in microsaccade rate (microsaccadic inhibition) and its associated microsaccade direction biasing was clearly dominated by the visual stimulus location, and this happened even despite each monkey’s individual prestimulus microsaccade direction biases.

**Figure 10.**
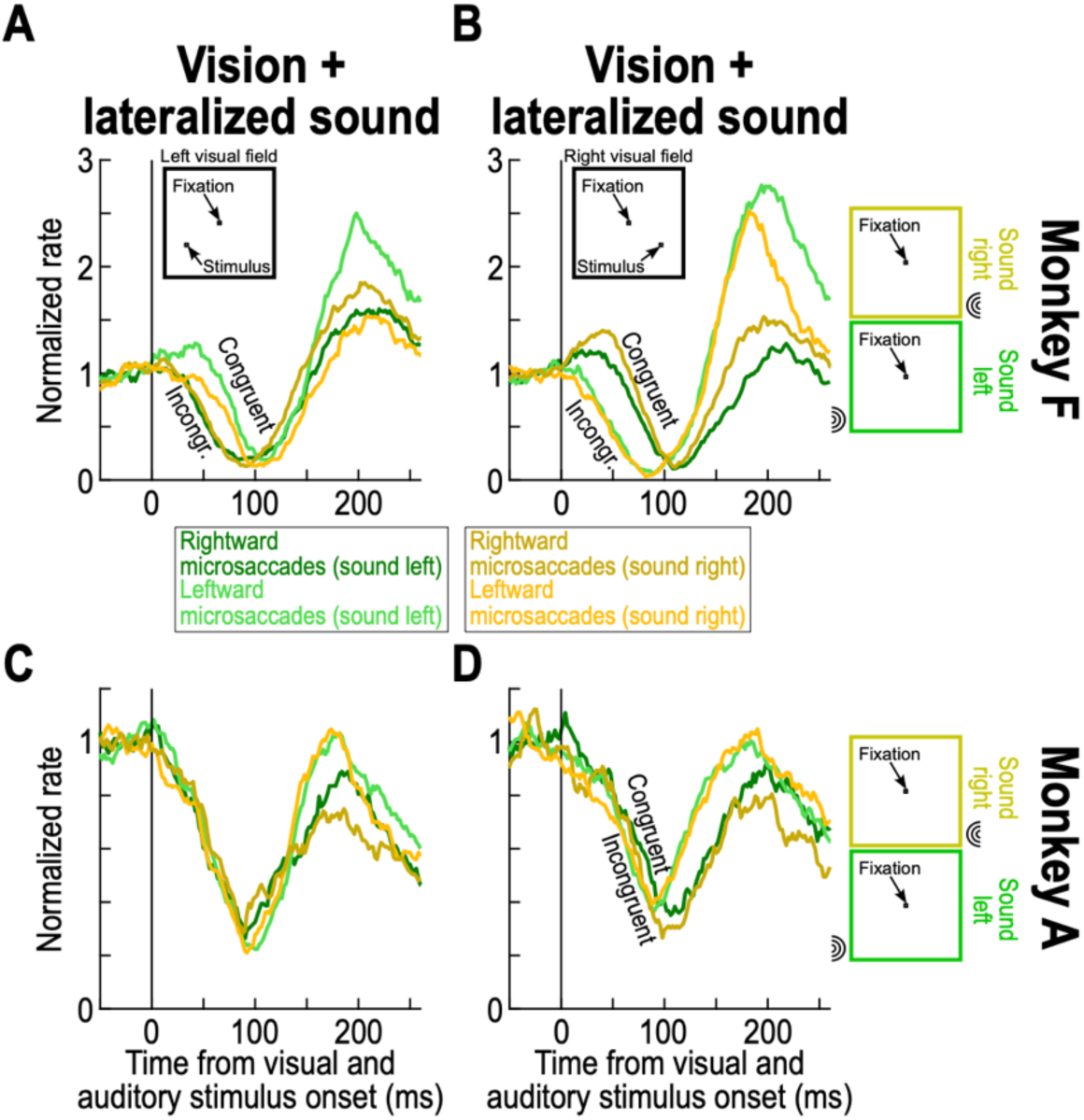
The results of Fig. 9, but after normalizing prestimulus rates. This figure shows the same data as in Fig. 9, but after normalizing to the prestimulus rate of every curve (Methods). This facilitates visualizing the directional modulation effects mentioned in the text and in Fig. 9.

Quantitatively, with a left sound, L_B25_ in monkey F was 44 ms and 76 ms for rightward and leftward microsaccades and a leftward visual stimulus, and it was 48 ms and 70 ms with a right sound (Fig. 9A). For the rightward visual field stimulus location (Fig. 9B), L_B25_ in the same monkey was 72 ms and 34 ms for rightward and leftward microsaccades and a left sound, and it was 76 ms and 32 ms with a right sound. These numbers, when comparing right and left sound groups to each other, were more similar to each other than the actual cueing effects expected from the visual stimulus location alone (Fig. 6), suggesting that the lateralized sounds barely influenced the visually-driven modulations. Statistically, with the visual stimulus on the left, the effect of the sound being on the left or right was not significant for leftward microsaccades (L_B25_ difference = 6 ms, *p_BH_* = 1, 95% CI [-16,16]), nor was it for rightward microsaccades (L_B25_ difference = -4 ms, *p_BH_* = 1, 95% CI [-12,12]). Similarly, with the rightward visual stimulus, the L_B25_ difference between the right and left sounds was not significant for either rightward (L_B25_ difference = 4 ms, *p_BH_* = 0.9385, 95% CI [-10,10]) or leftward (L_B25_ difference = -2 ms, *p_BH_* = 0.9698, 95% CI [-18,18]) microsaccades. Thus, the timing difference of L_B25_ was bigger when considering the congruency of microsaccade directions with the visual stimulus location than the congruency of microsaccade directions with the sound pulse location.

For monkey A and a left visual field visual stimulus (Fig. 9D), the L_B25_ difference between the leftward and rightward sounds was not significant for either leftward or rightward microsaccades (L_B25_ difference = 4 ms, *p_BH_* = 1, 95% CI [-18,18] and L_B25_ difference = -10 ms, *p_BH_* = 1, 95% CI [-30,32], respectively). Similarly, for the right visual field visual stimulus (Fig. 9E), the L_B25_ difference between the rightward and leftward sound conditions was not significant for either rightward or leftward microsaccades (L_B25_ difference = -12 ms, *p_BH_* = 0.8633, 95% CI [-20,18] and L_B25_ difference = -12 ms, *p_BH_* = 0.8633, 95% CI [-20,18], respectively). Thus, even with spatially-lateralized sounds, vision dominated the early cue-driven modulations of microsaccades in our monkeys.

There were other interesting features of the data in Fig. 9 that are worth documenting. First, a spatially-lateralized sound alone did not cause strong microsaccadic inhibition, consistent with all of our prior observations (with spatially lateralized or nonlateralized sounds). This can be seen in Fig. 9C, F for each monkey (compare to the strong microsaccadic inhibition in the vision plus sound trials, and also in the vision only trials of Fig. 6). Second, with a rightward sound alone, monkey A showed an early reduction in rightward microsaccades and a slightly smaller increase in leftward microsaccades at approximately 50 ms after sound onset (Fig. 9F; yellowish curves). This appeared as an early modulation in the microsaccade rate curves with a paired visual onset, especially in the right visual field visual stimulus condition (see Fig. 9E; yellowish curves). However, this early modulation effect in the vision plus sound trials was not enough to alter the fact that the visually-driven microsaccadic inhibition remained unaffected by the sound location (black oblique arrows in Fig. 9E illustrate this). Finally, after the microsaccadic inhibition phase (e.g. after more than 100 ms from sound pulse onset), monkey F showed a subtle increase in microsaccade rate on the sound-only trials (Fig. 9C). With a rightward sound, the increase happened for both rightward and leftward microsaccades equally (yellowish curves); with a leftward sound, the increase was stronger for leftward microsaccades (bright green curve). These so-called post-inhibition effects likely reflect top-down control over microsaccades (17), when recovering from the reflexive stimulus-induced inhibition (9, 17, 18, 27, 71), and they can explain why the yellowish and greenish curves in this monkey diverged from each other after the end of microsaccadic inhibition in the vision plus sound conditions of Fig. 9A, B.

In summary, all of our analyses revealed that vision dominated sound in driving the classic cue-driven modulations in microsaccade rate and directions in our rhesus macaque monkeys.

## Discussion

We were motivated by the question of how saccadic inhibition, and related ocular drift phenomena (10, 11), might be mediated in the brain. Clearly, sensory information needs to arrive at the final oculomotor control pathways with very short latencies, since measurable changes in eye movements can occur as early as 50-100 ms after sensory stimulus onset. However, this alone is not sufficient knowledge to pinpoint which sensory-motor pathways are involved in the inhibition and related phenomena, or which sensory-motor pathways are more prominent than others in these phenomena in the healthy brain.

Our starting point in the current study was a potential anatomical and physiological dissociation between the temporal and spatial aspects of visually-driven saccadic inhibition (17, 18, 20). By this, we mean that both the timing and metrics (e.g. spatial directions) of saccades can be affected during saccadic inhibition phases, and that these effects can be mediated by distinct brain circuits: one set of neurons might cancel ongoing saccade plans at the time of stimulus onset by raising the threshold for saccade triggering; and another set of neurons might bias the saccade metrics for the motor plans that do escape being canceled, by having visual bursts in topographic maps (1, 18, 22). If so, then adding a spatially-nonlateralized sound to a visual stimulus onset can alter the temporal component of saccadic inhibition, without altering the spatial biasing caused by the visual stimulus onset itself. And, adding a spatially-lateralized sound can alter both the temporal and spatial components.

In our first set of experiments, with spatially-nonlateralized sound pulses, we injected an additional “countermanding” command to visually-driven inhibition, without otherwise differentially influencing the spatial landscape of visual (or auditory) stimulus configurations. We found something intriguing: spatially-nonlateralized sounds alone did not cause substantial inhibition in comparison with the visual modality (Figs. 2, 3), but they still accelerated visually-driven inhibition (Fig. 3). Thus, we succeeded in modifying the temporal component of visually-driven saccadic inhibition. This observation is experimentally useful, because one can now use spatially-nonlateralized sounds as a way to speed up the temporal aspect of saccadic inhibition in the presence of visual stimuli, without the sounds alone themselves being as direct a driver for the observed eye movement modulations.

In our second set of experiments, we experimentally manipulated the spatial component of saccadic inhibition. Here, we placed the sound and visual stimulation sources either in spatial congruence or in spatial conflict with each other. Remarkably, we observed that the visual modality was the primary determinant of microsaccade direction modulations in our experiments.

The combined results from our experiments provide plenty of motivation for future neurophysiological investigations. For example, one hypothesis in the literature (1, 17, 18, 20) is that the temporal processes associated with saccadic inhibition can critically depend on omnipause neurons in the brainstem’s premotor oculomotor control network (72–75). These neurons gate saccade generation, but past work (primarily in cats) has shown that they also exhibit sensory responses (76–78); these responses functionally raise the threshold for saccade triggering, which in turn alters saccade rates. Importantly, the visual responses of these neurons are feature-tuned in monkeys (79, 80), in a manner that is consistent with the feature tuning properties of saccadic inhibition in these animals (24). Interestingly, we did not observe any evidence for adaptation of microsaccadic inhibition (Fig. 4), which might suggest that visual responses in omnipause neurons might not adapt to repeated sensory exposure, unlike visually-responsive neurons in many other brain areas. Similarly, on the side of saccade metrics, it has been suggested that the superior colliculus contributes to the spatial component of visually-driven saccadic inhibition (16, 17). Since both the superior colliculus and omnipause neurons are multisensory (41, 42, 46, 59, 78, 81–85), paradigms like ours can prove insightful about the pathways for inhibition. These paradigms also enable investigating other structures, such as the inferior colliculus (52, 86–88), which could (in the end) prove to be as critical as omnipause neurons and the superior colliculus. Such types of neurophysiological questions inspired by our behavioral results have the potential to allow substantial progress towards finally understanding much more of the neural substrates for saccadic inhibition, despite the phenomenon itself having been behaviorally documented for over two decades ago now.

Another intriguing observation that we made is that with spatially-nonlateralized sounds, there was an interference effect with the visually-driven direction modulations, especially in monkey F (Figs. 5, 6). On the one hand, this might be expected: a spatially-diffuse sound can introduce noise in the spatial representations encoding the appearing visual stimulus in the environment (42, 53, 54, 59, 69). On the other hand, with spatially-lateralized sounds, there was barely any alteration in the visually-driven direction modulations with a sound congruent or incongruent with the hemifield of the visual stimulus (Fig. 9). This is a somewhat surprising observation when compared to the spatially-nonlateralized sound situation, but it could also represent an opportunity for neurophysiological investigations, as mentioned above, since sound could be used as a modulatory input on an otherwise dominant visually-driven phenomenon. Indeed, we did see a clear and consistent impact of spatially-lateralized sounds on the temporal aspect of microsaccadic inhibition (Fig. 8), suggesting a boosting due to multisensory integration. It is also interesting that, in humans, it was suggested that spatial cueing by sounds did not cause systematic microsaccade direction modulations (89).

One additional possibility for explaining the above observations associated with spatially-lateralized sounds is that the auditory effect occurred earlier than the visual effect, resulting in the visual effect for spatially-lateralized sounds to be unaffected by spatial congruence with the visual stimulus. For example, in monkey A, especially for a rightward sound source, there was a slight and early biasing of microsaccades in the sound-only trials (Fig. 9F; yellowish curves). However, while this early biasing did cause an early biasing of the visually-driven rate curves in this monkey (Fig. 9D, E; yellowish curves), it does not really explain why the visually-driven inhibition timing was unaffected by the congruence of the sound location relative to the visual stimulus location. For example, the temporal aspects of microsaccadic inhibition (Fig. 8) were clearly influenced by the spatially-lateralized sounds (and not just in a very early phase after stimulus onset). A more likely possibility could be that the oculomotor system is better equipped to deal with competing lateralized representations of sensory stimuli (e.g. vision on one side and sound on another) than with a globally introduced noise injection into the system, like was the case with our spatially-nonlateralized sounds. Moreover, we believe that there was still some amount of interference by the spatially-lateralized sound trials when compared to the visual-only trials. For example, the inhibition times of the congruent and incongruent microsaccade rate curves were more different from each other in Fig. 6A (visual-only trials) than in Fig. 9A (vision plus spatially-lateralized sound trials). Thus, there was still likely some interference by the spatially-lateralized sound, but the key result remains that microsaccade direction modulations did not depend on whether the sound was congruent or incongruent with the visual stimulus location.

In all, our results revealed that vision does dominate sound in classic cue-driven modulations in microsaccadic eye movements in macaque monkeys. This does not mean that there cannot be any saccadic inhibition with auditory stimuli. In contrast, there is plenty of evidence for this in humans. For example, single auditory pulses, whether unilateral or bilateral, reliably induce microsaccadic inhibition in humans, although its strength and temporal dynamics tend to differ from those elicited by visual stimuli (7, 34, 55, 56, 68). Microsaccadic inhibition was also observed in selective attention paradigms, even in response to distractor stimuli, although it was weaker in that case (90). Furthermore, microsaccadic inhibition was observed in oddball sound paradigms, specifically in response to rare, oddball sound transients (71, 91–93). In fact, in our own results, monkey A showed mild inhibition (and more than monkey F) during the spatially-nonlateralized sound condition (Fig. 3). While this might reflect the higher sound intensities used for this particular monkey (Methods), we think that monkey F was actually the more sensitive monkey to the sounds, and that is why we reduced the intensities for this animal (Methods). Therefore, it remains to be seen whether sound intensity systematically causes saccadic inhibition in monkeys under sound-only conditions or not.

Finally, so far, we have always put forward the interpretation that our results indicate that when a visual stimulus is paired with sound, then the visual modality dominates the observed effects on the oculomotor system. Of course, stimulus onsets, in general, can reflexively capture attention (94, 95), and we know that microsaccadic inhibition is a component of a robust correlation between microsaccades and attention shifts (64, 65). Thus, our results inform effects of multisensory integration on reorienting of attention. For example, if the microsaccadic modulations are sped up by multisensory stimulation like we found, then it is likely that dynamics of attentional orienting might also do so. In our case, we were primarily focused in our interpretations on highlighting the more low-level mechanisms (such as visual and auditory bursts in sensory and sensory-motor structures throughout the brain) because this can help explicitly inform further neurophysiological investigations exploring the neural substrates of saccadic inhibition and related phenomena.

## Author contributions

Conceived and designed research: TM, MPB, ZMH

Analyzed data: TM, ZMH

Performed experiments: TM, MPB, YY, ZMH

Interpreted results of experiments: TM, MPB, YY, ZMH

Prepared figures: TM, ZMH

Drafted manuscript: TM, ZMH

Edited and revised manuscript: TM, MPB, YY, ZMH

Approved final version: TM, MPB, YY, ZMH

